# Chemical genetic screening identifies nalacin as an inhibitor of GH3 amido synthetase for auxin conjugation

**DOI:** 10.1101/2022.05.29.493926

**Authors:** Yinpeng Xie, Ying Zhu, Na Wang, Min Luo, Tsuyoshi Ohta, Ruipan Guo, Zongjun Yu, Yalikunjiang Aizezi, Linlin Zhang, Yan Yan, Yujie Zhang, Hongyu Bao, Yichuan Wang, Ziqiang Zhu, Ancheng Huang, Yunde Zhao, Tadao Asami, Hongda Huang, Hongwei Guo, Kai Jiang

## Abstract

Root system is critical for plant growth and development. To develop plant growth regulators functioning in root development, we performed a phenotype-based chemical screen in Arabidopsis and identified a chemical, nalacin, that mimicked the effects of auxin on root development. Genetic, pharmacological and biochemical approaches demonstrated that nalacin exerts its auxin-like activities by inhibiting indole-3-acetic acid (IAA) conjugation that is mediated by Gretchen Hagen 3 (GH3) acyl acid amido synthetases. The crystal structure of Arabidopsis GH3.6 in complex with D4 (a derivative of nalacin) together with docking simulation analysis revealed the molecular basis of the inhibition of group II GH3 by nalacin. Sequence alignment analysis indicated broad bioactivities of nalacin and D4 as inhibitors of GH3s in vascular plants, which were confirmed, at least, in tomato and rice. In summary, our work identifies nalacin as a potent inhibitor of IAA conjugation mediated by group II GH3 that plays versatile roles in hormone-regulated plant development and has potential applications in both basic research and agriculture.

## Introduction

In vascular plants, the root system is an important organ for the communications between the soil and the plant body (1). During the evolution, a flexible root system provides adaptive responses to internal signals and external environmental stimuli, and realizes the fitness and adaptation of plants (2). Indole-3-acetic acid (IAA), the main natural auxin, regulates plant root development on the basis of its local concentration (1, 3). Thus, IAA homeostasis must be tightly controlled by coordination of its biosynthesis, transport, storage and degradation (4).

IAA biosynthesis pathways have been well studied (4–6). The indole-3-pyruvate (IPyA) pathway, the prevailing route for IAA biosynthesis, consists of a two-step conversion of tryptophan (Trp) to IAA through the successive actions of TRYPTOPHAN AMINOTRANSFERASE OF ARABIDOPSIS1/TRYPTOPHAN AMINOTRANSFERASE RELATEDs (TAA1/TARs) and YUCCA flavin monooxygenases (5). Arabidopsis (*Arabidopsis thaliana*) with loss of function in TAA1/TARs or YUCCAs show dramatic defects in root, floral and vascular system development (7, 8).

Similar with auxin biosynthesis, auxin inactivation is also important for plant growth and development. Recent studies showed that IAA oxidation and conjugation are the two major routes for auxin inactivation (9). DIOXYGENASE FOR AUXIN OXIDATION (DAO), which belongs to the 2-oxoglutarate-dependent Fe(II) dioxygenase family, was first identified as an enzyme for auxin oxidation in rice (*Oryza sativa*) (10). A rice *dao* mutant showed elevated IAA levels and decreased production of 2-oxoindole-3-acetic acid (oxIAA), the major catabolic product of IAA, in the anthers and ovaries (10). Two DAO homologs, DAO1 and DAO2, have been identified in Arabidopsis, and their roles in auxin oxidation were characterized (11, 12). A *dao1* mutant with substantially decreased oxIAA levels showed no obvious changes in IAA levels but had an increase in IAA-Glu, indicating the importance of amino acid conjugation to the regulation of IAA homeostasis (11, 12). Recently, Hayashi et al. showed that DAO1 accepts IAA-Asp/Glu conjugates as substrates and catalyzes their oxidation (13).

The Gretchen Hagen 3 (GH3) proteins are a family of acyl-acid-amido synthetases that conjugate amino acids to various substrates (14, 15). According to the sequence homology and substrate preference, nineteen *GH3* genes in Arabidopsis can be categorized into three groups (16). Group II GH3, including GH3.1, GH3.2, GH3.3, GH3.4, GH3.5, GH3.6, GH3.9 and GH3.17, has been reported to mainly catalyze IAA conjugation to amino acids as a reversible stock (13, 15). In addition to IAA, GH3.5 displays wide preference for substrates, including 1-Naphthaleneacetic acid (NAA), phenylacetic acid (PAA), salicylic acid (SA), and benzoic acid (BA) (17). GH3.11 from group I prefers to catalyze conjugation of jasmonic acid (JA) to isoleucine, leading to its activation (14, 18). GH3.12 and GH3.7 from group III accept isochorismate as substrate to produce isochorismate-glutamate, a precursor that can be converted to SA (19, 20). GH3.15, as a member of group III, prefers to conjugate amino acids to indole-butyric acid (IBA), 2,4-dichlorophenoxylacetic acid (2,4-D) and 4-(2,4-dichlorophenoxy) butyric acid (2,4-DB) (21, 22). In contrast to the well-defined biochemical activities as mentioned above, the studies on biological roles of *GH3* genes including those in charge of IAA conjugation *in planta* lag behind due to their functional redundancy.

Chemical biology provides a powerful toolset of small molecules with which to explore biological processes (23). Small organic molecules can be applied to any plant tissue at appropriate concentrations at any stage of the plant life cycle, and provide the advantage of avoiding complications arising from gene redundancy and lethal mutations (24). Chemical genetics is widely applied in the study of IAA biosynthesis, signaling and transport, and many compounds have thereby been identified as regulators in these processes (25). Although many chemical regulators have been developed and widely used for regulating auxin biosynthesis, transport and signaling (25, 26), the regulators of auxin catabolism pathway are still rare. AIEP, a rationally designed inhibitor of GH3, displays potent inhibition on IAA conjugation *in vitro* (27), despite that its biological effects in broader plant species are yet to be defined. A recent preprint reports KKI as a GH3 inhibitor, which has been applied to the study of the physiological roles of GH3s (13, 28).

Here, we performed a phenotype-based chemical screen for root development in Arabidopsis and identified nalacin that displays auxin-like activities. Based on *in vitro* and *in planta* assays, we confirmed that nalacin exerts its auxin-like bioactivities by directly binding to several group II GH3 enzymes and inhibiting their catalysis of IAA conjugation. Crystal structural analysis together with docking simulation provides the molecular basis for the preference of nalacin as an inhibitor across group II GH3 of Arabidopsis and indicates its broad bioactivity in other vascular plants, which has been demonstrated in tomato and rice. In summary, we report nalacin and its derivative D4 as inhibitors of GH3 enzymes that catalyze amido conjugation of IAA, and provide a chemical tool for the functional study of *GH3* genes in hormone-regulated plant development as well as potential agricultural application.

## Results

### Nalacin is a root architecture regulator with auxin-like activities

To develop plant growth regulators (PGRs) for root architecture regulation, we performed a chemical screen (using a building block containing 11,800 chemicals from Life Chemicals) by observing the morphological changes in the roots of Arabidopsis wild-type Col-0 (Figure 1A). We discovered that the compound *N*-[4-[[6-(1*H*-pyrazol-1-yl)-3-pyridazinyl]amino]phenyl]-3-(trifluoromethyl)benza mide (Figure 1B) has versatile effects on the development of primary root, adventitious root and root hair together with other effects on hypocotyl elongation and cotyledon epinasty (Figure 1C), which are reminiscent of the effects of auxin. We designated the candidate compound as nalacin for its function as a <u>n</u>on-<u>a</u>uxin-scaffold-<u>l</u>ike <u>a</u>uxin <u>c</u>onjugation <u>in</u>hibitor, as indicated by the results detailed later in this article. Further physiological analyses showed that nalacin inhibited primary root growth and promoted adventitious root formation, lateral root number and root hair elongation in a similar manner to IAA (Figure 1D–F and Figure S1). Consistent with these auxin-like activities, we found that nalacin triggered fluorescence signal changes in *DII-VENUS* and *DR5::GFP*, two marker lines for auxin response (29, 30), in the same manner as IAA (Figure 2A, B), suggesting that nalacin plays a role in activating auxin signaling.

**Figure 1.**
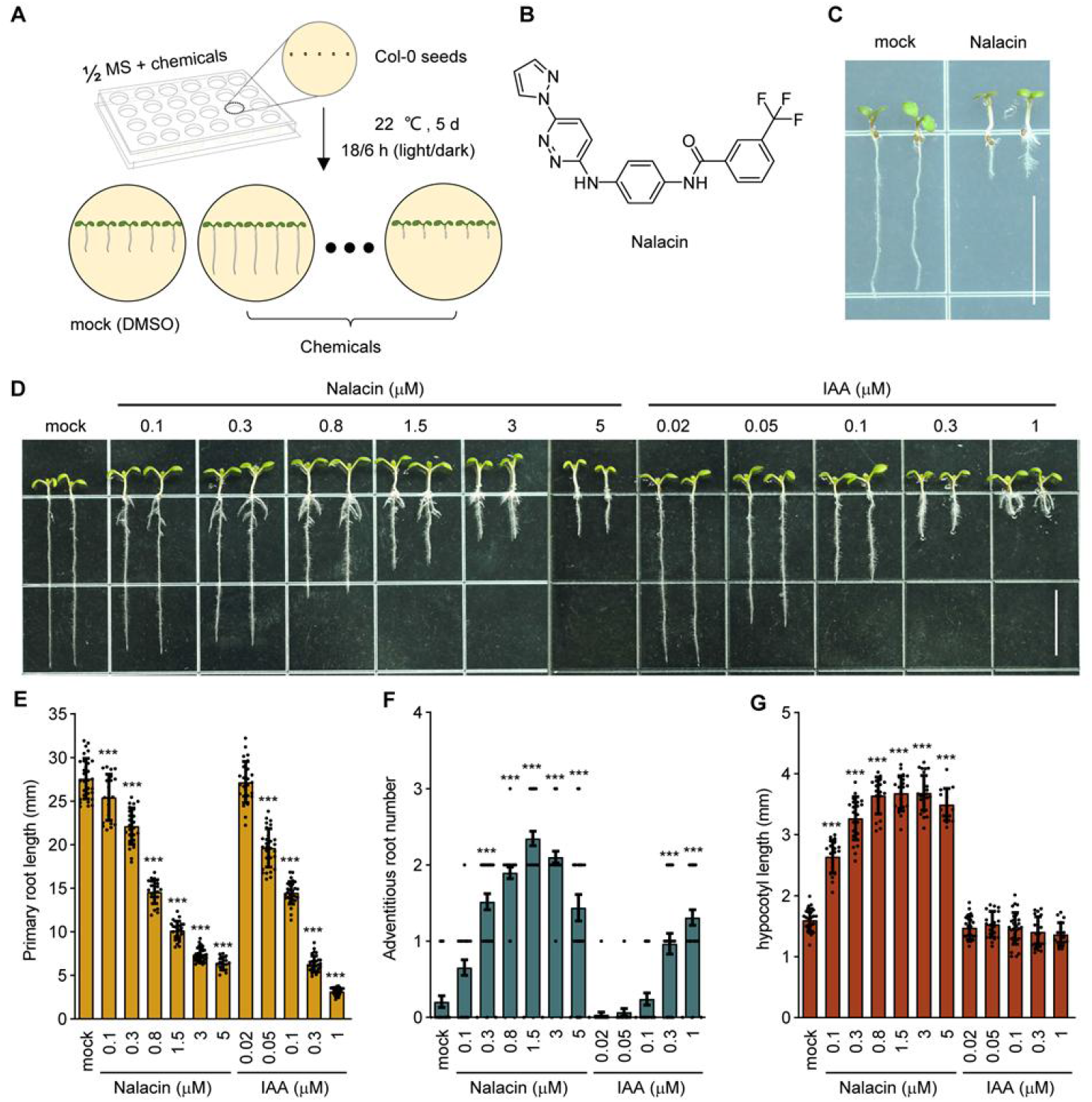
Nalacin partially mimics IAA in inducing root phenotypes. (**A**) A schematic graph of chemical screening with 11,800 small chemicals for plant growth regulator of root development. (**B**) The chemical structure of N-[4-[[6-(1H-Pyrazol-1-yl)-3-pyridazinyl]amino]phenyl]-3-(trifluoromethyl)benzamide (CAS registry number 1019105-44-22), which is designated as nalacin. (**C**) 5-d-old Col-0 seedlings grown vertically on MS medium containing 10 μM nalacin or DMSO as the mock control. Scale bar = 10 mm. (**D**) 6-d-old Col-0 seedlings grown vertically on MS medium containing gradient concentrations of nalacin, IAA, or DMSO as the mock control. Scale bar= 10 mm. (**E-G**) Quantification of the primary root length, adventitious root number and hypocotyl length of seedlings shown in (**D**). Values represent means and ±SD (n = 15). Statistical significances were analyzed by two-way ANOVA along with Tukey’s comparison test (\*\*\**P* < 0.001). Data in (**D-G**) were derived from experiments that were performed three times with similar results, and representative data from one replicate were shown.

**Figure 2.**
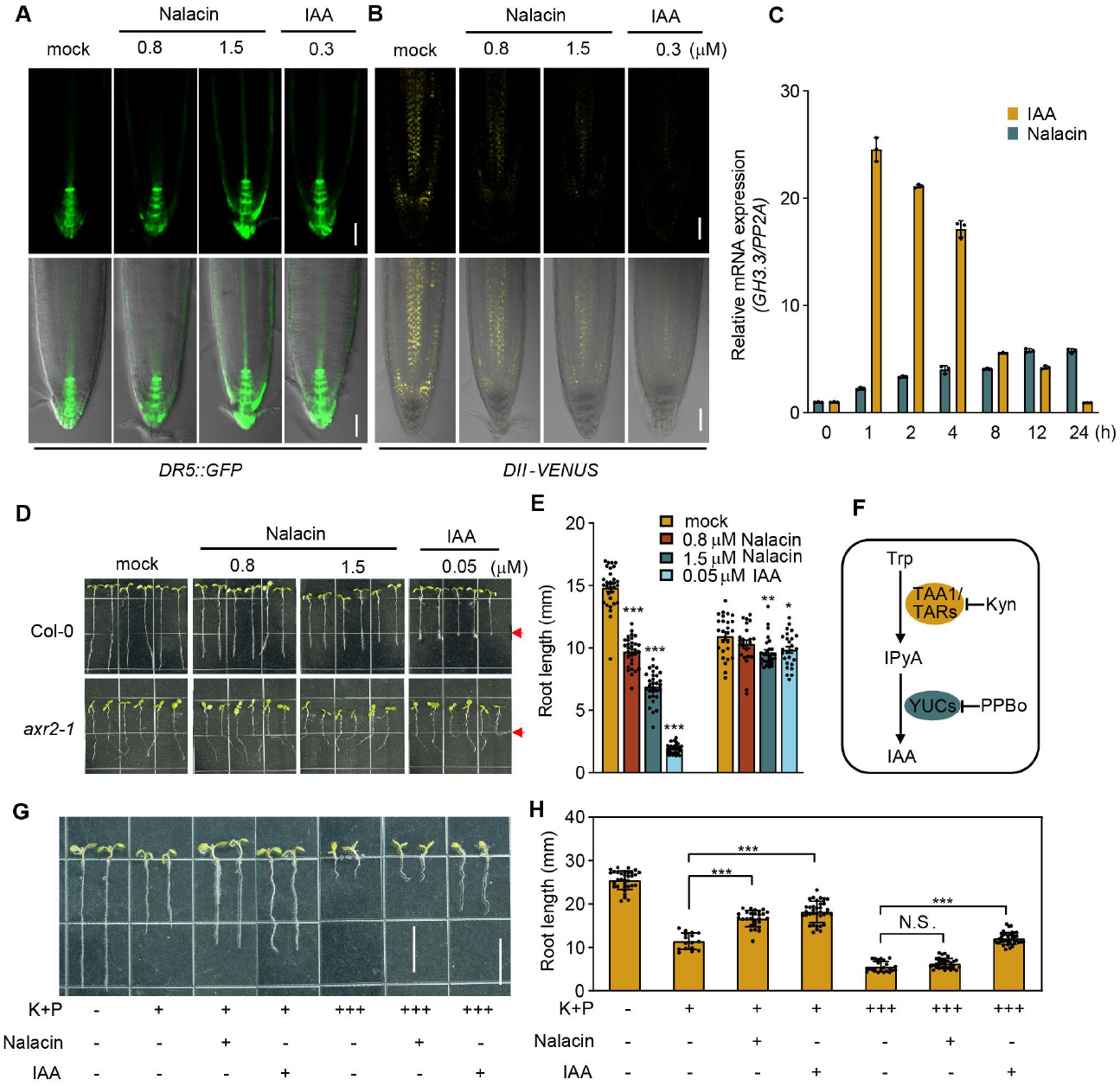
Nalacin activates auxin signaling in IAA biosynthesis-dependent manner. (**A**) Observation on the GFP fluorescence in the primary root of *DR5::GFP* reporter line. 4-d-old seedlings were transferred to MS medium containing nalacin, IAA, or DMSO for 24 h treatment before observing. Scale bar = 50 μm. (**B**) Observation on the YFP fluorescence in the primary root of *DII-VENUS* line. 4-d-old seedlings were transferred to MS medium containing nalacin, IAA, or DMSO for 24 h treatment. Scale bar = 50 μm. (**C**) Time-course expression of *GH3.3* in 6-d-old seedlings treated with 3 μM nalacin or 0.3 μM IAA in liquid MS medium. (**D**) 6-d-old Col-0 and *axr2-1* seedlings grown vertically on MS medium containing various concentrations of nalacin, IAA, or DMSO as the mock control. Scale bar = 10 mm. (**E**) Quantification of the primary root length of seedlings shown in (**D**). (**F**) A schematic diagram of Trp-dependent pathway for IAA biosynthesis. Trp, tryptophan; IPyA, indole-3-pyruvic acid; IAA, indole-3-acetic acid. TAA/TARs, TRYPTOPHAN AMINOTRANSFERASE OF ARABIDOPSIS1/TRYPTOPHAN AMINOTRANSFERASE RELATEDs; YUCs, YUCCA flavin monooxygenases; Kyn, L-kynurenine, an inhibitor of TAA/TARs; PPBo, 4-phenoxyphenylboronic acid, an inhibitor of YUCs. (**G**) 6-d-old Col-0 seedlings grown on MS medium containing combinatory treatment of 0.5 mM (+) or 10 mM (+++) Kyn and PPBo (K+P) together with nalacin (1.5 μM), IAA (0.02 μM), or DMSO. Scale bar = 10 mm. (**H**) Quantification of the root length shown in (**G**). Values represent means and ±SD (n ≥ 15). Statistical significances were analyzed by two-way ANOVA along with Tukey’s comparison test (\**P* < 0.05, \*\**P* < 0.01, \*\*\**P* < 0.001). Each dataset was derived from experiments that were performed three times with similar results, and representative data from one replicate were shown.

We also noticed additional and stronger effects of nalacin in promoting adventitious root formation and hypocotyl elongation in comparison with IAA (Figure 1D, F, G). By observing the β-glucuronidase (GUS) staining in a *DR5::GUS* reporter line, we found that nalacin triggered *DR5*-driven GUS signal in basal hypocotyl and lateral root primordium, which was not detected in the IAA-treated seedlings (Figure S2). *GH3.3*, a marker gene for auxin rapid response, is rapidly upregulated by auxin treatment, and then decreases due to negative feedback regulation (31, 32). We found that, compared to IAA, nalacin produced a slower, longer-lasting induction of *GH3.3* expression (Figure 2C). Together, these data indicate that nalacin is a novel regulator of root architecture that may act by activating auxin signaling in a spatiotemporal manner different from that of IAA.

### Nalacin suppresses root growth in an IPyA-dependent auxin biosynthesis pathway

To gain insights into nalacin’s mode of action, we investigated its relationship with auxin via genetic and pharmacological analyses. *axr2-1* is a gain-of-function mutant line with a non-degradable IAA7 that disrupts an early step in auxin response (33, 34). Both IAA and nalacin largely reduce their inhibition on primary root growth in *axr2-1* mutant (Figure 2D, E), suggesting that a functional auxin signaling pathway is required for their bioactivity. L-kynurenine (Kyn) and 4-phenoxyphenylboronic acid (PPBo) are two inhibitors of IAA biosynthesis that target the TAA1 and YUCs enzymes, respectively, to decrease endogenous IAA levels (Figure 2F) (35, 36). Combination treatment with Kyn and PPBo (K+P) resulted in a short primary root (Figure 2G, H) due to their pharmacological role in decreasing local IAA level. Treatment with either nalacin or IAA partially restored the primary root growth that was suppressed by 0.5 μM K+P (Figure 2G, H). However, when suppressed by a higher concentration of K+P (10 μM), the primary root growth could be partially restored by treatment with IAA but not nalacin (Figure 2G, H). Collectively, these results suggest that nalacin is not an auxin mimic and that its auxin-like activity is dependent on endogenous IAA biosynthesis, possibly via its impact on IAA homeostasis.

### Nalacin increases endogenous IAA level via inhibition of IAA conjugation

To validate the effects of nalacin on IAA homeostasis, we quantified the levels of IAA, its biosynthesis precursors (Trp and IPyA) and major inactivated metabolites in Col-0, including oxIAA and amino-acid-conjugated IAAs (IAA-Glu and IAA-Asp) whose formation are catalyzed by DAOs and GH3s, respectively (11, 12, 15). Compared to the mock treatment, the Trp and IPyA contents were not altered after nalacin treatment (Figure 3A, B), while nalacin increased IAA levels and decreased the levels of IAA-Asp and IAA-Glu without changing the levels of oxIAA (Figure 3C-F). These results strongly imply that nalacin exerts its auxin-like activities via inhibiting the inactivation of IAA by amino acid conjugation, which leads to higher levels of active IAA.

**Figure 3.**
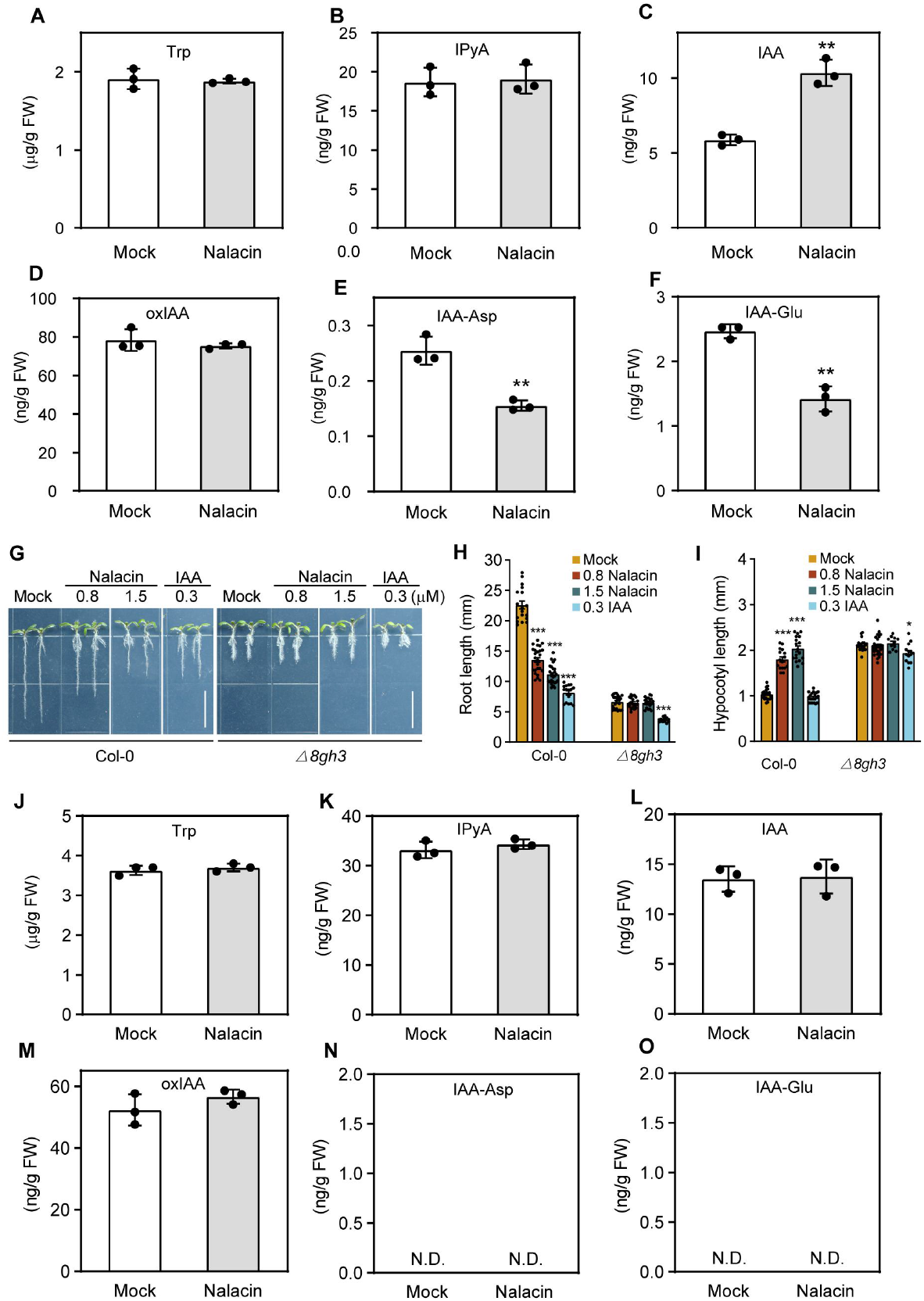
The *gh3* octuple mutant is resistant to nalacin. (**A-F**) Determination of IAA, IAA biosynthetic precursors (Trp and IPyA) and major metabolic products (IAA-Glu, IAA-Asp, and oxIAA) under nalacin or DMSO treatment in Col-0. Trp, Tryptophan; IPyA, indole-3-pyruvic acid; IAA, indole-3-acetic acid; oxIAA, 2-oxoindole-3-acetic acid; IAA-Asp, IAA-asparate; IAA-Glu, IAA-glutamate. DAOs, DAO1 and DAO2; GH3s, group II GH3s. (**G**) Representative seedlings of 6-d-old Col-0 and *gh3* octuple mutant (*△8gh3*) grown on MS medium containing nalacin, IAA, and DMSO. (**H-I**) Quantification of the root length and hypocotyl length shown in (**G**). Values represent means and ±SD (n ≥ 15). (**J-O**) Determination of IAA, IAA biosynthetic precursors (Trp and IPyA) and major metabolic products (IAA-Glu, IAA-Asp, and oxIAA) in *△8gh3* mutant. Statistical significances were analyzed by two-way ANOVA along with Tukey’s comparison test for (**H-I**) and student’s t-test for (**A-F, J-O**) (\**P* < 0.05, \*\**P* < 0.01, \*\*\**P* < 0.001). Each dataset was derived from experiments that were performed three times with similar results, and representative data from one replicate were shown.

As stated above, nineteen *GH3* genes can be divided into three phylogenetic groups (Figure S3A) (15, 16), among which the group II GH3 is responsible for amino acid conjugation of IAA. A recent report showed that a mutant with loss of function of all eight group II *GH3* (*△8gh3*) displayed a short primary root and an elevated number of adventitious roots (37), a pattern reminiscent of the effects of nalacin (Figure 1C). We found that *△8gh3* was resistant to nalacin treatment, while still being responsive to exogenous IAA treatment (Figure 3G–I). Moreover, the endogenous levels of IAA and its precursors and metabolites showed no response to nalacin treatment in the octuple mutant (Figure 3J–O). These results indicate that nalacin could target group II GH3 to influence IAA homeostasis.

### Nalacin directly interacts with GH3s and suppresses their enzymatic activity on amino acid conjugation of IAA

To test whether group II GH3 enzymes are the direct targets of nalacin, we selected GH3.3, GH3.6 and GH3.17 as representatives for each subgroup of group II GH3 based on phylogenetic analysis (Figure S3B), and produced recombinant proteins of these three GH3 proteins via *Escherichia coli*. We used a biolayer interferometry (BLI) assay, an optical technique to measure protein–small molecule interactions (38), to analyze the interaction of nalacin for GH3 proteins. We found that nalacin displayed slow association and dissociation kinetics for GH3.3, GH3.6 and GH3.17 (*K*_d_ values of 51.6 μM, 140 μM and 28.5 μM, respectively) (Figure 4A–C), indicating a direct binding of nalacin to these GH3s. We further performed *in vitro* enzyme kinetic analyses, and found that nalacin inhibited the amino acid conjugation of IAA catalyzed by GH3.3, GH3.6, and GH3.17 at sub-micromolar (*K*_i_ values were 0.86 μM, 0.19 μM, and 0.4 μM respectively) (Figure 4D–F). The double-reciprocal plots of initial velocities (Lineweaver–Burk plots) indicate that nalacin is a competitive and uncompetitive mixed-type inhibitor, which could bind to both free enzyme and substrate-bound enzyme. Compared to AIEP, a competitive GH3 inhibitor with IAA scaffold (17, 27), nalacin functions as a <u>n</u>on-<u>a</u>uxin-scaffold-<u>l</u>ike <u>a</u>uxin <u>c</u>onjugation <u>in</u>hibitor with different inhibitory mode of action. It is worth to note that GH3.5 displays dual function in amino acid-conjugation of both IAA and SA (17, 39). Compared with the mock treatment, the endogenous concentration of SA in Col-0 was not altered upon nalacin treatment (Figure S4A). This result indicates that GH3.5-mediated SA conjugation is not affected by nalacin, which could be attributed to lower efficiency of GH3.5 in catalyzing SA (*Km* = 1171 μM) than that of IAA (*Km* = 45 μM) (39).

**Figure 4.**
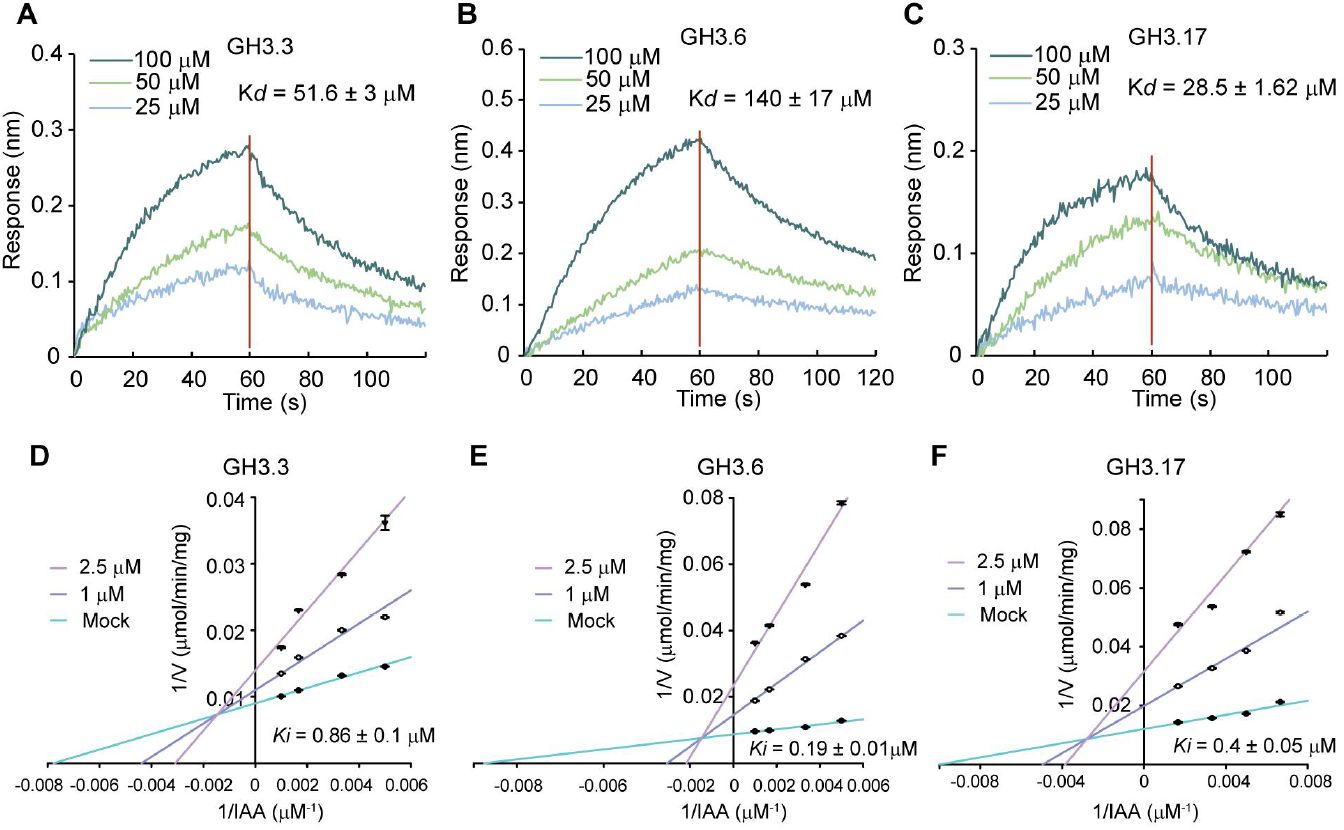
Nalacin interacts with GH3 proteins and inhibits the enzymatic activities of GH3s *in vitro*. (**A-C**) Biolayer interferometry assay on the kinetic interactions of nalacin with GH3.3, GH3.6 and GH3.17. Biotinylated GH3 proteins were loaded on SSA sensors followed by exposure to gradient concentrations of nalacin (25-100 μM). (**D-F**) Kinetic analysis of the inhibition of nalacin on GH3.3, GH3.6 and GH3.17. Activity assays were performed at a fixed aspartate (10 mM, for GH3.3 and GH3.6) or glutamate (10 mM, for GH3.17) with varying concentrations of IAA (0.15 mM-1 mM) in the absence or presence of 1 μM or 2.5 μM nalacin. The production of IAA-Asp (for GH3.3, GH3.6) and IAA-Glu (GH3.17) were detected by UPLC-MS. Double-reciprocal plots of initial velocities (Lineweaver-Burk plots) showing mixed-type inhibition. Values represent means and ±SD (n ≥ 3). Each dataset was derived from experiments that were performed three times with similar results, and representative data from one replicate were shown.

To investigate whether nalacin targets on group I and group III members, we selected GH3.11 and GH3.12 as representative members of group I and group III GH3 respectively. We observed no inhibitory effect of nalacin on GH3.12 for catalyzing its substrate 4-HBA into 4-HBA-Glu via *in vitro* assay (Figure S4C-E) (40). GH3.12 is also reported to catalyze the conjugation of glutamate to isochorismate and produce isochorismate-9-glutamate, which spontaneously converts into SA *in planta* (20). Consistent with our *in vitro* result, we observed no significant change of SA levels between nalacin-treated seedlings and the mock-treated ones in Col-0 and *△8gh3* (Figure S4A-B), indicating nalacin has no influence on GH3.12. In contrast, we found that nalacin suppressed the Ile-conjugation of JA catalyzed by GH3.11 with a mixed-type inhibition (Figure S5A). We also observed decrease in endogenous levels of JA-Ile in nalacin-treated Col-0 and *△8gh3* (Figure S5B-C). These results indicate that nalacin inhibits GH3.11 both *in vitro* and *in planta*. However, nalacin-induced phenotype changes in root and hypocotyl are not attributed to nalacin’s inhibition on GH3.11 as no significant difference was observed between Col-0 and *jar1-1*, a *gh3.11* mutant, in terms of these phenotypes, and their responses to nalacin were quite similar (Figure S5D-F). Thus, we focus on investigating nalacin’s mode of actions on group II GH3 enzymes that mediate auxin homeostasis and contribute to regulatory its roles in root development.

### Structure–activity relationship (SAR) relationship analysis of nalacin

During the nalacin synthesis process, we obtained four intermediate chemicals that partially share common moieties with nalacin. We designated them D1-D4 and tested their bioactivities (Figure S6). We observed no phenotypic changes in seedlings treated by D1 or D2. D3 displayed weak bioactivities in promoting adventitious root and inhibiting primary root growth at relatively high concentration level (1.5 μM) (Figure S6B-D). D4 showed an effect similar to nalacin in promoting adventitious root number and hypocotyl elongation, except that its inhibition of primary root growth was weaker (Figure S6B-E). In support of the physiological observations, we observed that D4 inhibited IAA-Asp production catalyzed by GH3.6 with a *K*_i_ of 0.3 μM (Figure S7). These results indicate that the *N*-(4-aminophenyl)-3-(trifluoromethyl)benzamide moiety represented by D3, rather than those represented by D1 or D2, is the basic structure for the bioactivities. Introducing either pyrazol-1-yl-pyrizadine or chloropyrizadine moieties into D3, which produces nalacin and D4, respectively, dramatically increased the bioactivity. These SAR studies identify a novel GH3 inhibitor D4, and provide information for further chemical modifications on nalacin.

### Structural analysis of the GH3.6-AMP-D4/nalacin complex

To investigate the molecular basis of nalacin and D4 inhibition of group II GH3 proteins, we tried to crystallize the Arabidopsis GH3.6 protein in complex with nalacin or D4 alone or in combination with other substrates or products, including ATP, ATPγS, AMP and Mg^2+^. After extensive trials, we successfully crystallized the GH3.6-AMP-D4 complex and solved its crystal structure to 2.40 Å (Supplementary Table 1). The structure of this complex adopts a closed active site conformation (Figure 5A) (17, 41), and the overall fold of GH3.6 resembles the fold of other GH3-family proteins, with a large N-terminal α/βfold domain (residues 1-446 in GH3.6), a small C-terminal domain (residues 460–612) and a hinge loop (residues 447–459) connecting them (Figure 5A, B) (17). AMP is bound in the nucleotide-binding site of GH3.6, which is conserved in other GH3s (17, 41), indicative of their universal enzymatic mechanism (Figure 5C). D4 is buried in a relatively hydrophobic channel that is also the acyl acid-binding pocket of the GH3-family proteins (17, 41), and has extensive interactions with GH3.6 and AMP (Figure 5A, B, D and Figure S8). For analysis, the structure of D4 can be subdivided into a trifluoromethylbenzamide portion (I), a phenyl portion (II) and an aminochloropyridazine portion (III) (Figure S6A). The interactions within part I are as follows (Figure 5B, D): the trifluoromethyl group establishes van der Waals contacts (distance ≤4.0 Å) and an F°°°H-P hydrogen bond with the phosphate group of AMP, with the latter implying that the phosphate group of AMP is protonated; the trifluoromethyl group forms contacts with Ala339, Ser340 and S341 in the β8–turn–β9 motif; the phenyl ring forms CH-π stacking with Met337 and Ala339 on one side and Ile312 on the opposite side, and also contacts Leu175 and the adenine moiety of AMP; and the carbonyl group is hydrogen-bonded to Tyr134. Within part II (Figure 5D), the phenyl group establishes T-shaped π-π stacking with Tyr134; and one edge of the phenyl group forms OH-π and CH-π attractions with Tyr179 and Val231, respectively. Lastly, within part III (Figure 5D), the amine group and one nitrogen atom of the pyridazine group are each hydrogen-bonded to Tyr195; the pyridazine group establishes Y-shaped π-π stacking with Tyr134 and Phe184, and also contacts Tyr178 and Thr193; and the chloride atom forms some contacts with Tyr178.

**Figure 5.**
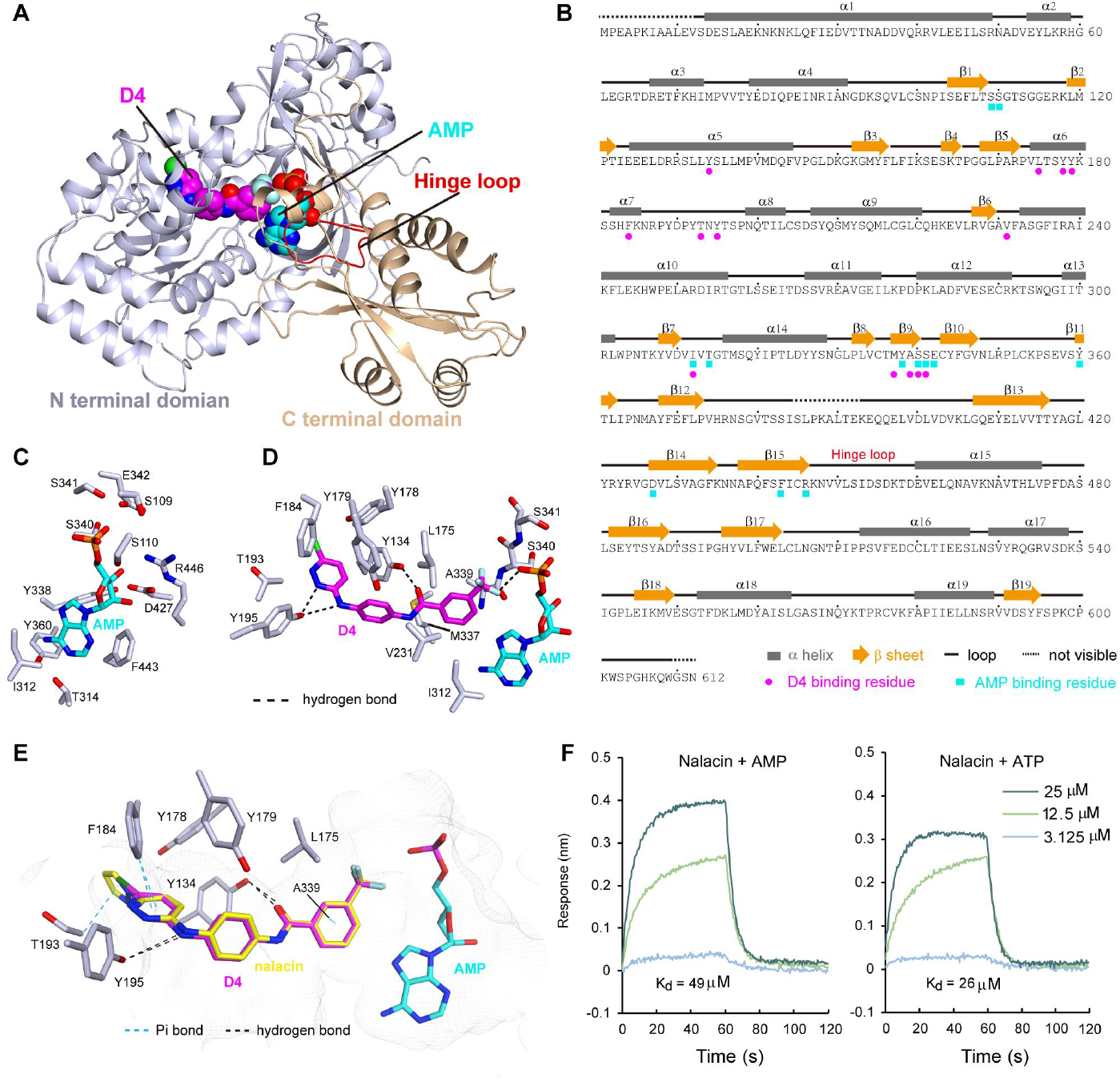
Structural analysis of the GH3.6-AMP-D4/nalacin complex. (**A**) Overall structure of the GH3.6-AMP-D4 complex, adopting a close active site conformation. The N- and C-terminal domains and the hinge loop connecting them are highlighted. The bound AMP and D4 molecules are shown in sphere mode. (**B**) The sequence and secondary structures of GH3.6 derived from the complex structure. The residues interacting with D4 and AMP are highlighted with magenta circles and cyan squares, respectively. (**C**) The conserved AMP-binding site of GH3.6. (**D**) An enlarged view showing the detailed interactions of D4 with GH3.6 and AMP. (**E**) A site view of molecular docked nalacin (in yellow color) superposed with D4 (in purple color) from the crystal structure. (**F**) Biolayer interferometry assay on the kinetic interactions of nalacin with GH3.6 in the presence of AMP or ATP. Biotinylated GH3 proteins were loaded on SSA sensors followed by exposure to gradient concentrations of nalacin (3.125-25 μM) and fixed concentration of AMP (100 μM) or ATP (100 μM). Data in (**F**) were derived from experiments that were performed two times with similar results, and representative data from one replicate were shown.

To analyze the molecular modes of action of nalacin, we performed a molecular docking simulation based on the structure of the GH3.6–AMP–D4 complex. As expected, we redocked D4 to GH3.6 in almost the same pose as observed in the crystal structure (Figure S9) and with a reasonable docking scoring (*S*–8.0900), suggesting the reliability of our docking process. Nalacin was docked to GH3.6 and superposed well with D4 except for its additional pyrazol moiety (Figure 6E), which introduces a new CH-π interaction with Thr193 and may contribute to its better affinity for GH3.6 (S –8.6907) and greater bioactivity as compared to D4 (Figure S6B–E). Compared to AIEP, another GH3 inhibitor (17, 27), nalacin occupies a broader space in the pocket of GH3 and possesses a pyrazol-1-yl-pyrizadine moiety that faces away from the enzymatic reaction center (Figure S10). D1, D2 and D3 have interaction poses that clearly differ from those of nalacin and D4, according to the docking analysis, and have low affinities for GH3.6, as represented by poor docking scorings (S –5.4018, –5.8495 and –6.5732, respectively) (Figure S9), in line with their bioactivities (Figure S6B–E). In view of the fact that ATP shares similar binding sites with AMP (17, 41) and nalacin contacts only the alpha phosphate group of AMP (Figure 5D), nalacin may also bind to GH3.6 in complex with ATP. We performed a BLI assay and confirmed that nalacin displayed fast association and dissociation kinetics with reasonable affinities for GH3.6 in the presence of AMP or ATP (*K*_d_ values of 49 and 26 μM, respectively) (Figure 5F). These biochemical results, together with the structural information, suggest that nalacin can inhibit AMP- or ATP-bound GH3s.

**Figure 6.**
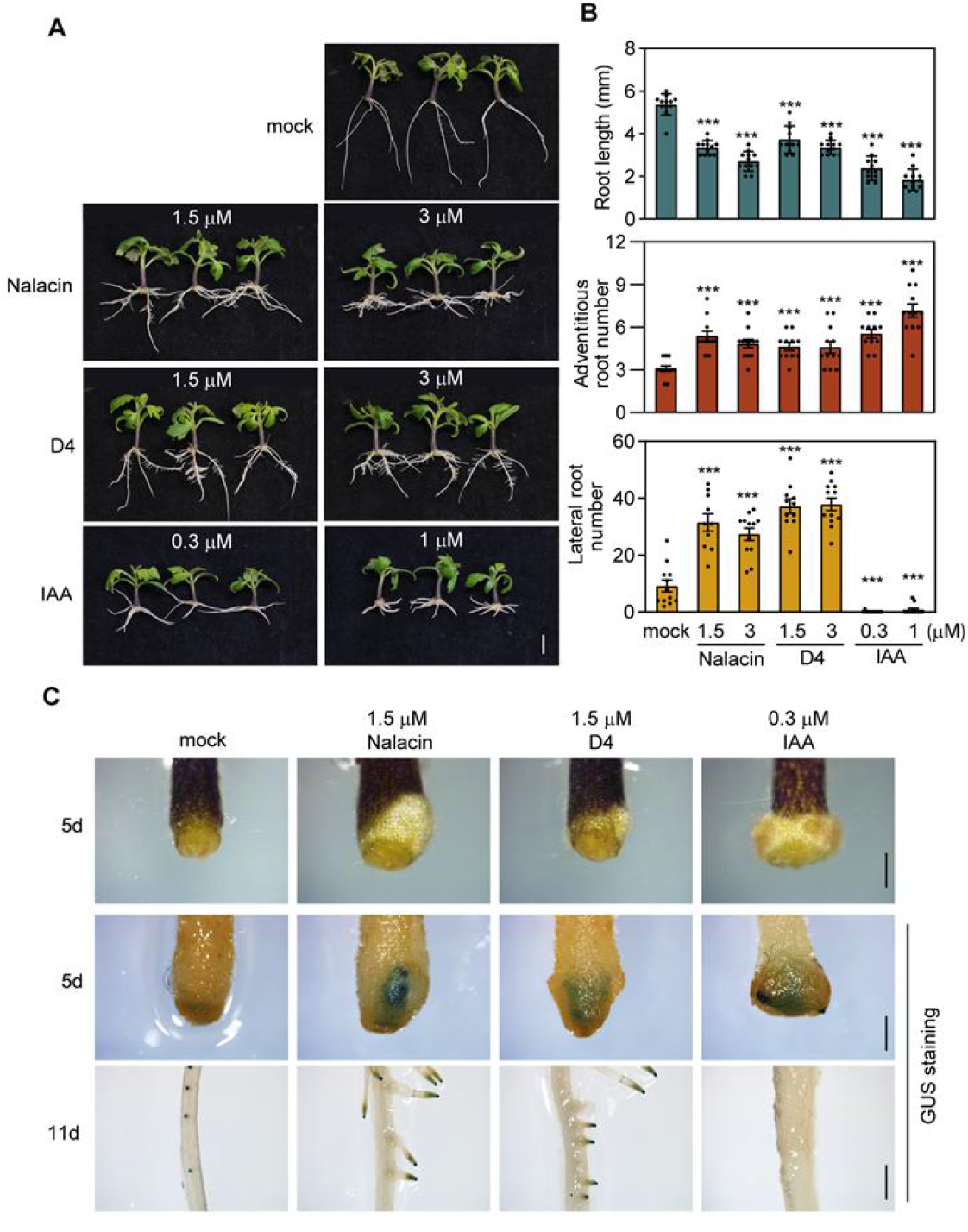
Nalacin and D4 promote adventitious root development in tomato. (**A**) Cutting propagation assay with tomato. 7-day-old tomato seedlings were detached with root and transferred to MS medium containing nalacin, D4, IAA or DMSO followed by additional cultivation for 10 days. Scale bar = 1 cm. (**B**) Quantification of the root length, adventitious root number and lateral root number shown in (**A**). (**C**) 7-day-old *DR5::GUS* seedlings of tomato with depletion of the roots were transferred to MS medium containing nalacin, D4, IAA, or DMSO followed by additional cultivation for several days. The GUS staining were performed on day 5 (cutting edge) or day 11 (regenerated roots) after transferring. Scale bar = 1 mm. Statistical significance was analyzed by two-way ANOVA along with Tukey’s comparison test (\*\*\**P* < 0.001). Data in (**A**-**C**) were derived from experiments that were performed two times with similar results, and representative data from one replicate were shown.

### 3D-structure-based analysis of nalacin’s mode of action in Arabidopsis and other plant species

Arabidopsis GH3 proteins share the same Ping-Pong reaction mechanism for catalyzing the conjugation of versatile substrates, including IAA, jasmonic acid and salicylic acid (17, 41, 42). Thus, whether nalacin is a selective inhibitor for a unique group of GH3s or a pan-inhibitor for all GH3s remained unknown. We analyzed the binding pockets of all GH3s by sequence alignment and by comparing their three-dimensional structures, obtained from the Protein Data Bank (PDB), AlphaFold (43) and our *de novo* homology modeling results (Figure S11). We found that the overall folds are generally conserved in each GH3 group (Figure S11A). As expected, all GH3s have a set of conserved residues for binding ATP and AMP (Figure S11B, D), which are the common substrate and product for catalysis by GH3s. In contrast, the residues constituting the pocket that binds nalacin/D4 are highly conserved in group II but not in groups I or III GH3s (Figure S11C, D), indicating a preference of nalacin/D4 for group II GH3s. It is noteworthy that 13 residues of GH3.6 for interacting with nalacin/D4 (indicated by round dots and diamond shapes in Figure S11D) are not all conserved in GH3.4 and GH3.9. Two residues Tyr134 and Thr193 in GH3.6 that form multiple interactions with nalacin/D4 were replaced by Gly and Ser respectively in GH3.4, while Tyr195 and Ile312 in GH3.6 were replaced by Leu and Val respectively in GH3.9 (Figure S11D), possibly leading to decreases in the binding affinities of nalacin/D4. In addition, we also noticed that GH3.10 and GH3.11 from group I, and GH3.7, GH3.12 and GH3.19 from group III possess several nalacin/D4-interacting residues (although not as many as in group II GH3s) (Figure S11D). These results and our *in vitro* and *in vivo* assays for the inhibitory effect of nalacin on GH3.11 and GH3.12 (Figure S4-5) indicate that nalacin could bind to and inhibit other GH3s with a different manner from that of group II GH3 members. In spite of those possibilities, our structural analyses together with biochemical and biological studies strongly suggest that nalacin and D4 exert their auxin-like bioactivities via preferably targeting group II GH3.

Previous studies indicated that GH3 proteins were conserved in the plant kingdom (16). We performed BLAST analysis and alignment of the homologs of Arabidopsis GH3.6, using them as representatives of group II GH3s from lower plants to higher plants, including moss, liverworts, fern, monocots and eudicots (Figure S12A, B). We found that the residues that consist of the pocket and the nalacin- and D4-interacting residues are highly conserved in vascular plant species, implying that nalacin and D4 should also have bioactivity in other higher plant species. To validate this hypothesis, we performed a tomato cutting propagation assay, in which auxin plays a pivotal role (44). We found that nalacin and D4 significantly induce adventitious root formation in a similar manner to IAA (Figure S13A–C). After long-time cultivation, nalacin and D4, but not IAA, triggered the initiation of lateral roots in adventitious roots (Figure 6A, B), a favorable phenotype in agriculture. We believe that this may be attributable to a longer-lasting effect of nalacin and D4 compared to IAA because GUS activity was observed only in the lateral root of *DR5::GUS* seedlings treated with nalacin and D4 (Figure 6C). Moreover, nalacin and D4 inhibited the root growth of rice seedlings (Figure S14) similarly to IAA (45). Taken together, the above results confirm that nalacin and its derivative D4 can inhibit group II GH3 and thus IAA conjugation, and have broad activities in various plant species.

## Discussion

Plants rely on the homeostasis of auxin to regulate plant growth and development, as well as to response to various environmental cues (46). Thus, it is critical for plants to fine-tune the levels of free IAA spatiotemporally, and auxin catabolism pathway is indispensable to this process. *GH3* was firstly identified as an early auxin responsive gene in soybean (*Glycine max*) (47). Subsequent studies have shown that the *GH3s* are spatiotemporally induced by many other plant hormones and ambient signals (such as biotic and abiotic stresses), and widely distribute in dicots and monocots (48), indicating their involvement in the plant development and environmental accommodation. Substantial genome analyses in various plants reveal great numbers of *GH3* genes and their high degree of redundancy that have been well illustrated by a series of studies in Arabidopsis (11, 15, 37). To overcome the genetic redundancy of *GH3*s, Guo *et al*. utilized the CRISPR/Cas9 gene editing method to generate *gh3* mutants with loss of function of multiple group II GH3s in Arabidopsis and confirmed their redundant functions in regulating root growth and a critical role of GH3.9 in fertility (37). *gh3oct* mutant was also reported to have enhanced tolerance to salinity and water deficit (49). However, a constitutive loss of function in *GH3*s results in infertility that limit the studies on GH3s with a spatiotemporal manner. In addition, the biological studies of GH3s in broader plant species still lag behind due to the time-consuming approach for generating mutants in multiple genes and/or technical challenges in genetic transformation in many non-model plants.

Chemical genetics is a powerful tool for studying plant biology, which can overcome gene redundancy and lethal mutation. By a phenotype-based chemical screening, we identified nalacin, an artificial small molecule that regulates root architecture in a similar manner to IAA (Figure. 1, 2A–C and Figure S1). Nalacin can directly bind to several group II GH3s (Figure 4A–C) and inhibit GH3-mediated conjugation of Glu or Asp to IAA (Figure 4D–F), leading to an increase in endogenous IAA levels *in planta* (Figure 3C). In support of that, nalacin-treated Col-0 mimicked the phenotypes of *gh3* octuple mutant (Figure 3G–I), which could not be observed in *gh3.1,2,3,4,6* (11) due to the redundant roles of *GH3.17* (37). In addition, *gh3* octuple mutant showed resistance to nalacin treatment in both phenotypic changes and perturbation of endogenous IAA levels (Figure 3L). These results demonstrate that nalacin can inhibit group II GH3s and overcome their redundant roles in catalyzing IAA conjugation. Compared to *gh3* octuple mutant with defects in whole development stage, nalacin is expected to be temporally applied to dissect the biological roles of GH3s at any time point in the whole lifespan of plants.

The expression of several *GH3* genes (*GH3.1, GH3.3, GH3.5* and *GH3.6*) is upregulated by auxin; while the GH3 enzymes act to inactivate intracellular auxin, which forms a negative feedback loop (16). This was also observed in our study, where exogenous IAA treatment caused a rapid increase followed by a rapid decrease in *GH3.3* expression (Figure 2C). However, nalacin treatment caused a mild, long-lasting increase in auxin signal (Figure 2C). In accordance with these modes of action, we observed GUS signal accumulation in hypocotyl and hypocotyl-root junction in seedlings treated with nalacin rather than IAA (Figure S2), which correlates with the stronger physiological effects of nalacin in promoting adventitious root formation and hypocotyl elongation (Figure 1D, F, G). These inconsistencies in the plant phenotypic changes in response to treatments with nalacin and IAA result from their different modes of actions in perturbing the endogenous IAA homeostasis. Due to the feedback regulation of IAA by IAA-induced *GH3* transcript levels and subsequent inactivation by GH3s-mediated conjugation, exogenous IAA treatment exerts bioactivity under strict regulation of GH3s upon their spatial-temporal expression pattern (37, 50–55). In contrast, owing to the inhibitory effects on GH3s, nalacin can trigger the accumulation of auxin and exert its auxin-like bioactivity without being strictly gated by GH3s. Thus, nalacin displays different morphological effects from IAA in a tissue-specific manner of GH3s. What’s more, the fact that nalacin is a stable and effective inhibitor that can nicely overcome gene redundancy provides a strategy for spatiotemporal regulation of GH3 enzyme, which can be used for both scientific research and agricultural application in non-model plants, such as root regeneration as demonstrated in this work.

In addition to nalacin, we also developed its derivative D4 and succeeded in resolving a complex structure of GH3.6 and D4 (Figure 5A-D). Based on the information from crystal structure and docking simulation, we found that the nalacin- and D4-interacting residues are highly conserved in group II GH3s in Arabidopsis and various vascular plants (Figure S11D, Figure S12), indicating their general bioactivities via targeting group II GH3s and thus perturbing IAA levels across the plant species. In support of that, we have confirmed that nalacin and D4 display auxin-like activities, at least, in Arabidopsis, tomato and rice (Figure 1, 6, and Figure S13,14). We also noticed that several GH3s from group I and III share multiple residues for interacting with nalacin and D4 in GH3.6 (Figure S11D), suggesting their potentiality as the targets of nalacin and D4. This is supported by the observation that nalacin suppressed JA-Ile production catalyzed by GH3.11, a group I GH3 member, *in vitro* and *in vivo* (Figure S5A–C). However, nalacin showed no inhibitory effect on GH3.12, a group III GH3 member (Figure S4C–E). As the substrates of many GH3s are yet to be determined, thus, in comparison with *in vitro* test one by one, a metabolome analysis in future could give a comprehensive understanding of nalacin-triggered metabolite changes and the possible targets of nalacin. Nevertheless, our current results corroboratively demonstrate that nalacin and D4 are effective inhibitors of IAA-conjugating GH3s that can be applied to regulate auxin homeostasis.

GH3-mediated IAA conjugation utilizes a “Bi Uni Uni Bi Ping-Pong” reaction that includes two successive steps (42). First, GH3 binds ATP and IAA successively, and catalyzes the formation of IAA-AMP (as an intermediate) and pyrophosphate by an adenylation reaction. Second, an amino acid such as aspartate or glutamate binds to the reaction pocket of GH3 and forms an amide bond with IAA by nucleophilic attack, producing amino-acid-conjugated IAA and AMP (42). Our crystal structure and docking simulation results provide a structural basis for nalacin’s binding and inhibitory mechanism in the view of the Ping-Pong reaction (Figure 5A–E). We observed that nalacin/D4 occupies a broad space in the GH3 pocket, including the IAA-binding site, which corresponds to the observed mix-type inhibition of nalacin/D4 against IAA in the enzymatic assay (Figure 4E, Figure S7). The van der Waals contacts and hydrogen bonds observed between nalacin/D4 and AMP could contribute to their affinity for GH3s. In consistence, we confirmed that nalacin can bind to GH3.6 in complex with ATP *in vitro* and displayed higher affinity for ATP-bound GH3.6 (Figure 5F) than the apo GH3.6 (Figure 4B). We also noticed that ATP altered the association and dissociation pattern for nalacin binding to GH3.6 (Figure 4B, 5F). Taken together, we propose that nalacin prefers to bind to and inhibit group II GH3s in complex with ATP in the first step of ping-pong reaction. In view of that nalacin is a mixed-type inhibitor for substrate IAA (Figure 4D–F) and the complexity of the Ping-Pong reaction, it remains possibility that nalacin could also inhibit GH3s with additional mode of actions.

AIEP, a rationally designed analog of the intermediate in GH3-mediated IAA-conjugation, was reported to be a potent inhibitor of grape GH3 enzymes (27), while the biological activity of AIEP *in planta* has not determined yet. KKI is recently reported to be an inhibitor of GH3 for catalyzing IAA conjugation in a preprint (28). According to the enzymatic assays, AIEP and KKI share the same inhibitory manner (competitive inhibition on IAA binding) on GH3s that mediate IAA conjugation. However, whether these two chemicals inhibit other IAA-unrelated GH3s is yet to be determined. Nalacin with definitely different structure with AIEP and KKI occupies the binding sites for IAA and amino acid, and further penetrates into an inner-side pocket (Figure S10). The inner-side pocket occupied by the pyrazol-1-yl-pyrizadine moiety of nalacin is distal to the enzyme reaction center, providing a potential avenue for genetic engineering of GH3s. The engineered GH3 variants would be expected to tolerate nalacin binding without impairing the original enzyme activity. In view of the auxin-like herbicidal effects of nalacin at high concentrations, the combinations of nalacin and the nalacin-tolerant GH3 variants may provide new tool kits for the design of herbicide-tolerant crops or for other deliberate regulatory changes.

In summary, we have identified and characterized nalacin and its derivative D4 as inhibitors of GH3s in hormone-regulated plant development, providing chemical tools to facilitate the study on the biological roles of GH3s in various plants and to be applied as a PGR in agricultural production. Last but not least, we demonstrate a computer-aided pipeline for structural analysis and docking simulation, which can be utilized for assessing the potential binding affinity of nalacin for GH3s in other plant species. It is expected to develop novel GH3 inhibitors for various plant species based on the chemical scaffold of nalacin/D4 and the strategies set-up in this work.

## Materials and Methods

Detailed descriptions of plant materials and growth conditions, and treatments are described in *SI Materials and Methods*. Small molecule library and screen information, root hair length measurement and fluorescence observation, GUS staining, measurement of IPA, IAA and IAA catabolites, GH3 enzymatic activity assay and kinetic analysis, biolayer interferometry analysis of binding activity of nalacin with GH3 proteins, crystallization, X-ray data collection and structure determination, molecular docking simulation, Homology modelling of group III GH3s, and chemical preparation according to protocols described in *SI Materials and Methods*.

## Acknowledgements

This work was supported by the National Natural Science Foundation of China (Grant No. 31911540070 to HG and Grant No. 21907049 to KJ), the National Key Research and Development Program of China (Grant 2019YFA0903904 to HG), a JSPS Grant-in-Aid for Scientific Research (Grant 18H05266) to TA, the Key Laboratory of Molecular Design for Plant Cell Factory of Guangdong Higher Education Institutes (SUSTech) (2019KSYS006 to HG), the Guangdong Innovative and Entrepreneurial Research Team Program (Grant No. 2016ZT06S172) to KJ, the Shenzhen Science and Technology Program (Grant No. KYTDPT20181011104005 to KJ and Grant No. KQTD20190929173906742 to HG), and the China Postdoctoral Science Fund Project (Grant No. 2020M672406 to YX.). We thank Dr. Hua Li (SUSTech CRF), Dr. Lin Lin (SUSTech CRF) and Ms. Suying Gao (SUSTech) for the kind help of mass spectrometry analyses, Dr. Wei Huang for the support of tomato tissue culture, Dr. Dan Zhang and Mr. Yuping Qiu for the help of confocal laser scanning training, Dr. Yang Peng for the root hair analysis, Ms. Qiuhua Yang and Dr. Xing Wen for the support of protein purification. We also thank all members in Guo lab for stimulating discussion and suggestions. We thank the staffs from the BL17B1/BL18U1/BL19U1/BL19U2/BL01B beamlines of National Facility for Protein Science in Shanghai (NFPS) at Shanghai Synchrotron Radiation Facility, for assistance during data collection, and Dr. Yang Zhao (Shanghai Center for Plant Stress Biology, Chinese Academy of Sciences) for supporting the chemical library.

## Author Contributions

Y.X., H.G., and K.J. designed and guided the research. Y.X., Y.Z., N.W., Y.A., L.Z., Y.Y., Y-J.Z., Y-D.Z., A.H. and Y.W. performed research. Y.Z. and Z.Z. performed the chemical library screens and the initial characterization of small molecule candidates. T.O. and T.A. performed chemical synthesis. R.G. and Y.Z. generated the genetic material of *gh3* multiple mutants. N.W., M.L., H.B., and H.H. performed the crystal structure study. K.J. performed molecular docking simulation and homology modelling. Y.X., N.W, H.H., H.G., and K.J. wrote the paper.

## Availability

Atomic coordinates and structure factors for the reported crystal structure have been deposited in the Protein Data bank under accession numbers 7VKA (the AtGH3.6-AMP-D4 complex). Other data are available upon reasonable request.

---

**Figure S1.**
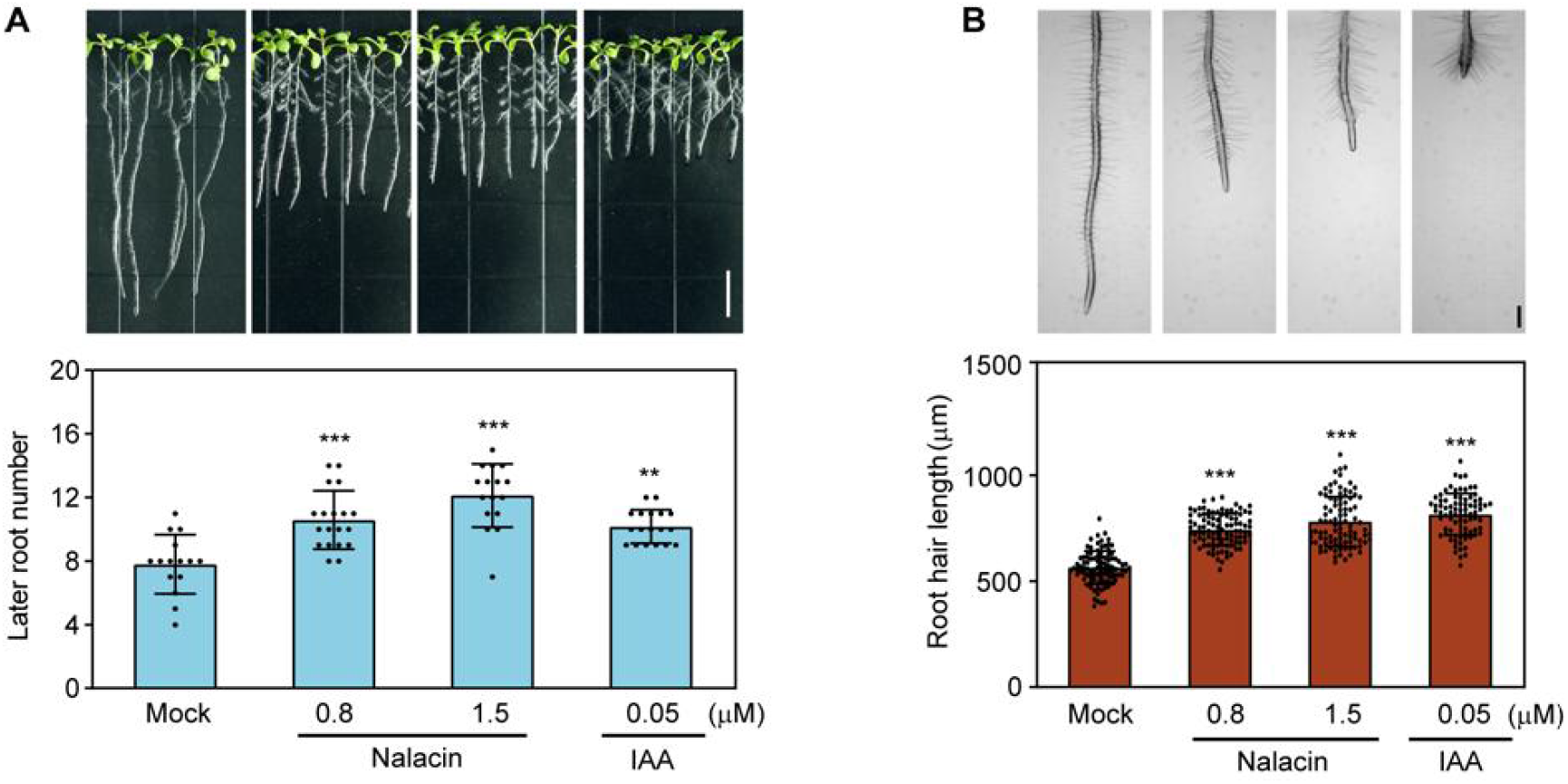
Nalacin promotes lateral root initiation and root hair elongation. (**A**) 4-day-old Col-0 seedlings grown on MS medium were transferred to MS medium containing nalacin, IAA followed by additional cultivation for 4 days. Top panel, representative seedlings treated by nalacin, IAA, and DMSO as the mock control. Scale bar = 10 mm. Bottom panel, quantification of the lateral root number shown in the top panel. Values represent means and ±SD (n ≥ 15). (**B**) 4-day-old Col-0 seedlings grown on MS medium were transferred to MS medium containing nalacin, IAA, or DMSO followed by additional cultivation for 24 hours. Top panel, representative root hairs treated by nalacin, IAA, and DMSO. Scale bar = 200 μm. Bottom panel, quantification of the root hair length shown in top panel. Values represent means and ± SD (n ≥ 80). Statistical significance was analyzed by two-way ANOVA along with Tukey’s comparison test (\*\**P* < 0.01; \*\*\**P* < 0.001).

**Figure S2.**
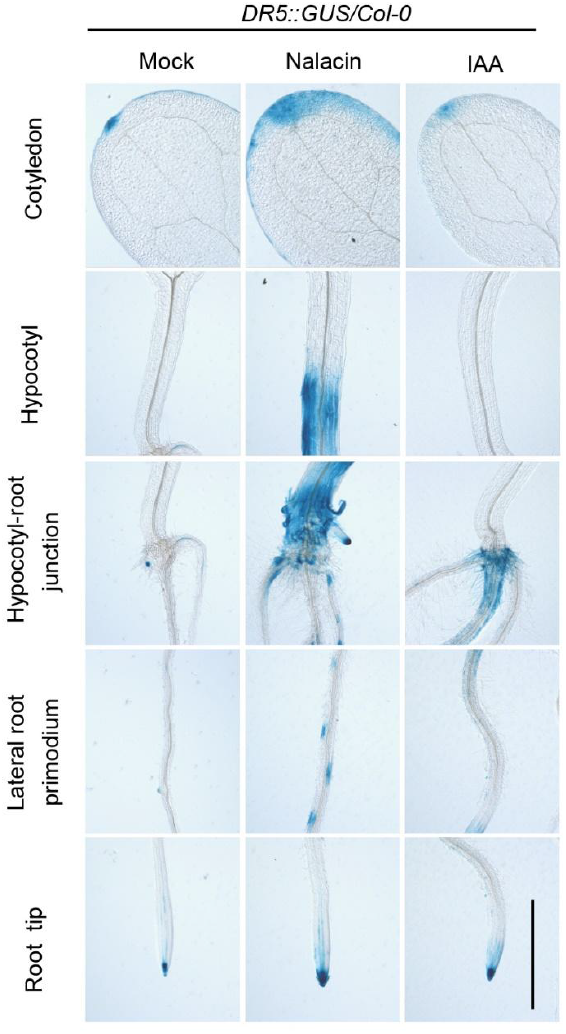
Nalacin activates auxin signaling in a tissue specific pattern. GUS staining observation in 6-day-old *DR5::GUS* seedlings that were grown on MS medium containing 1.5 μM nalacin, 1 μM IAA, or DMSO as the mock control. Scale bar = 100 mm.

**Figure S3.**
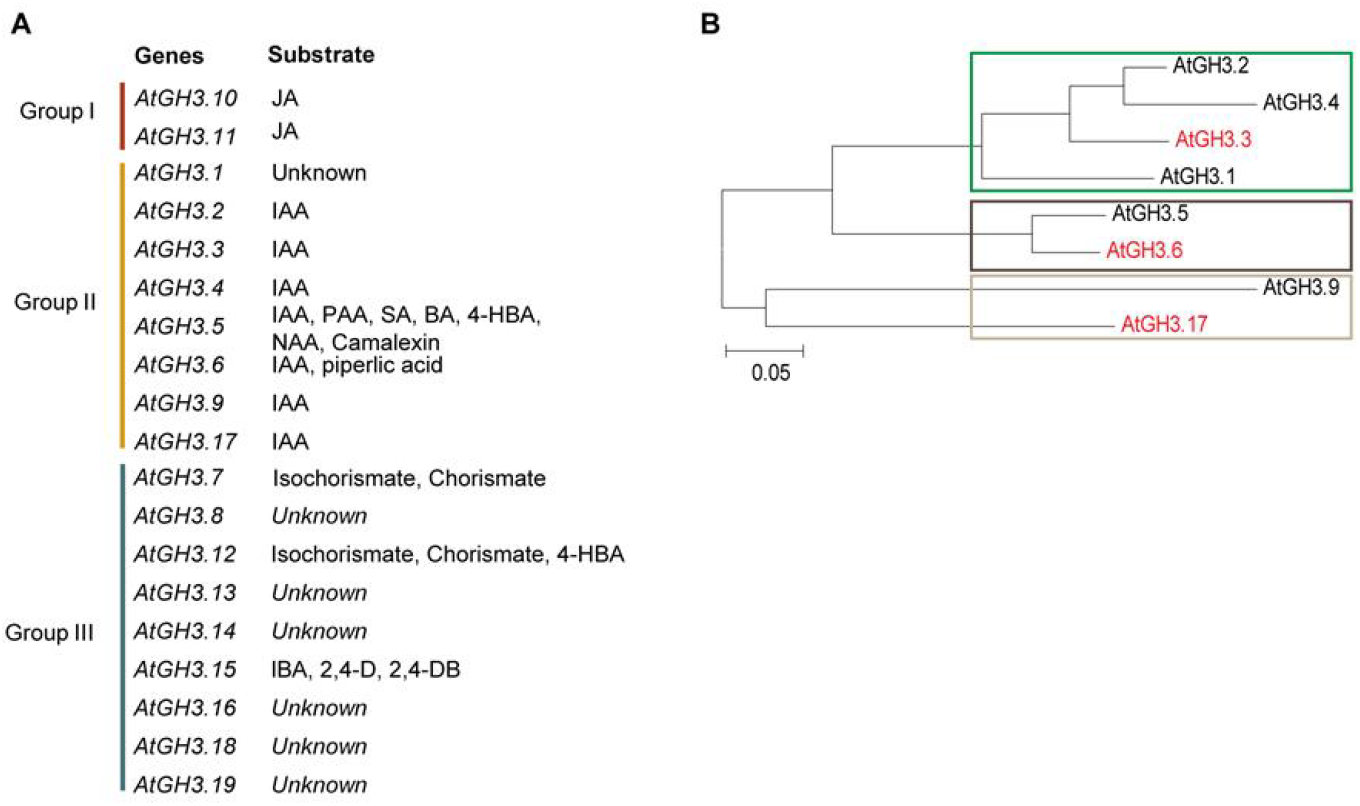
Phylogenetic analysis of GH3s proteins in Arabidopsis. (**A**) Possible substrates for each GH3. JA, jasmonic acid; IAA, indole-3-acetic acid; PAA, phenylacetic acid; SA, salicylic acid; BA, benzoic acid; 4-HBA, 4-hydroxybenzoic acid; NAA, 1-Naphthaleneacetic acid; IBA, indolebutyric acid; 2,4-D, 2,4-dichlorophenoxyacetic acid; 2,4-DB, Butyl 4-(2,4-dichlorophenoxy)butanoate. (**B**) The group II GH3 can be divided into three sub-groups indicated by rectangles. GH3.3, GH3.6, and GH3.17 are representatives of each subgroup for *in vitro* experiments.

**Figure S4.**
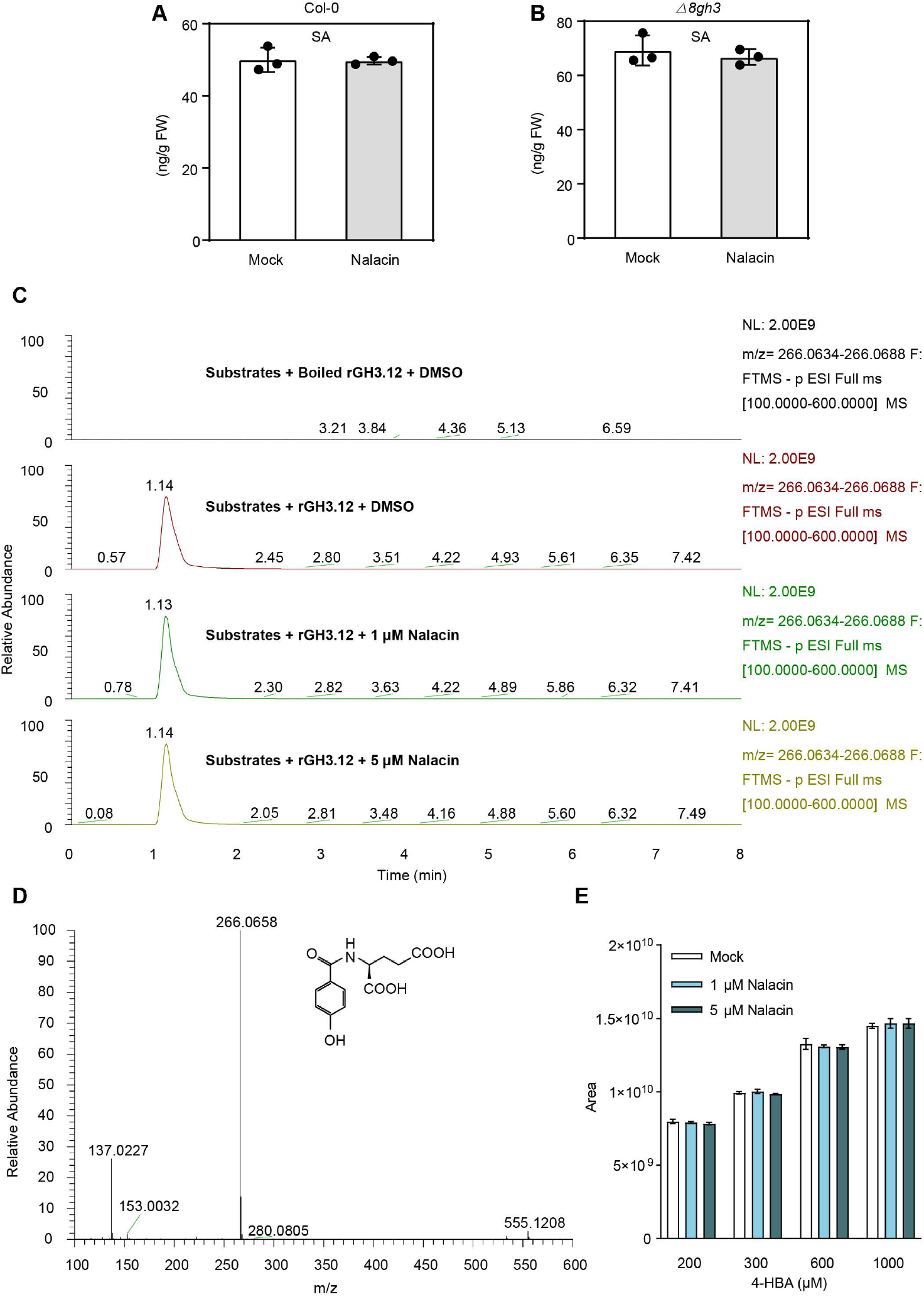
Nalacin has no effect on GH3.12 *in vitro* and *in vivo*. (**A-B**) Determination of SA in Col-0 and *Δ8gh3* under nalacin or DMSO treatment. (**C-E**) LC-MS/MS analysis of reaction products of 4-HBA-Glu catalyzed by GH3.12. Substrates (4-HBA, ATP, Ile) and recombinant protein of GH3.12 (rGH3.12) with or without addition of nalacin were incubated at 30 °C. Mass spectrum of the product of 4-HBA-Glu were showed in (**C**). (**D**) is Q-TOFMS/MS on m/z = 268.0658 with 4-HBA-Glu structure. The peak areas of 4-HBA-Glu under different concentration of 4-HBA and nalacin were shown in (**E**). Each data was derived from experiments that were performed three times with similar results, and representative data from one repetition were shown.

**Figure S5.**
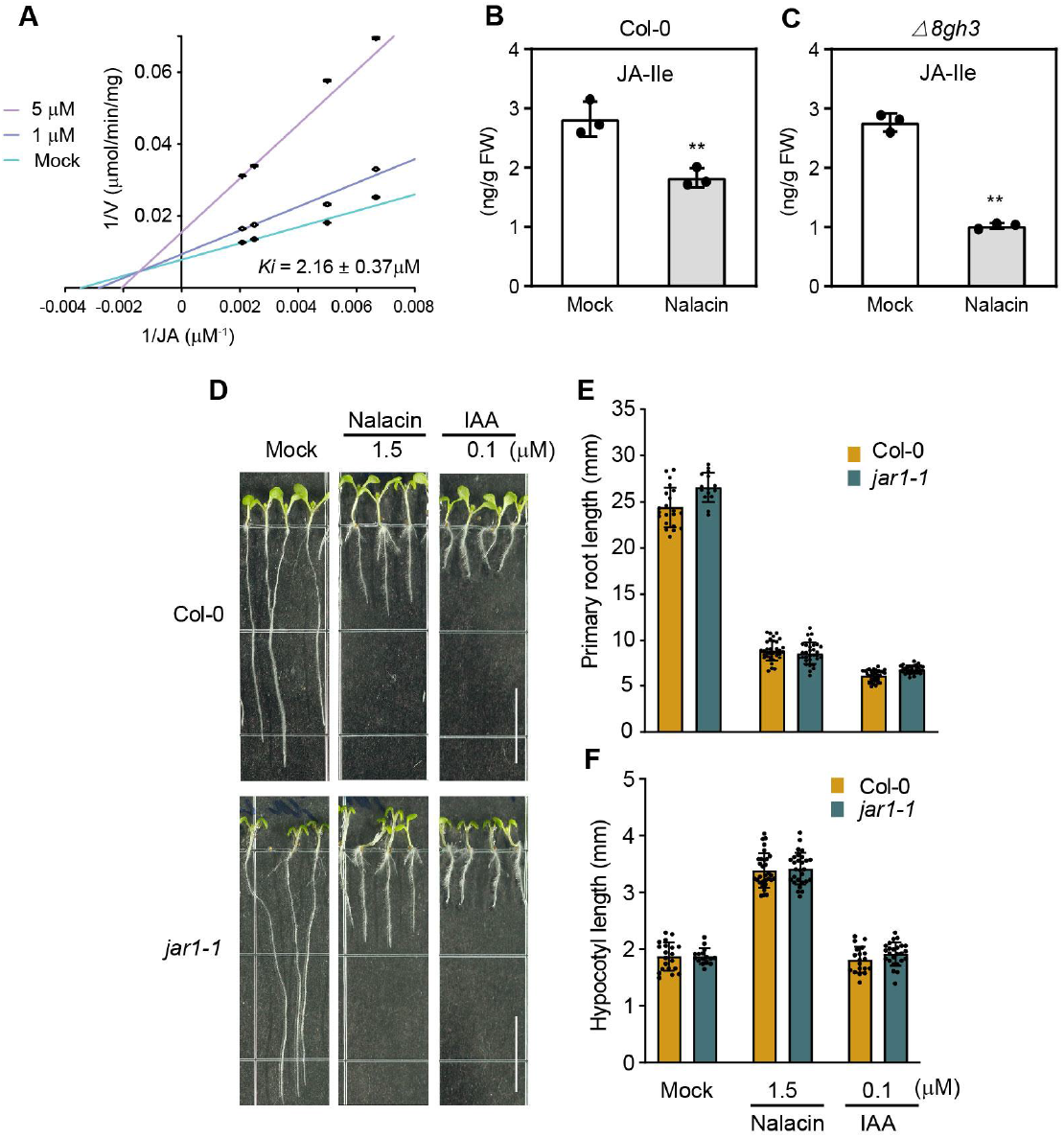
Nalacin inhibits enzyme activities of GH3.11. (**A**) Enzyme kinetics analysis of the inhibition of nalacin on GH3.11. Activity assays were performed at a fixed isoleucine (10 mM) with varying concentration of JA (0.1 mM-1 mM) in the absence and presence of 1 μM or 5 μM nalacin. The production of JA-Ile was detected by UPLC-MS. Double-reciprocal plots of initial velocities (Lineweaver-Burk plots) showing a mixed-type inhibition. Values represent means and ±SD (n ≥ 3). (**B-C**) Determination of JA-Ile in Col-0 and *Δ8gh3* under nalacin or DMSO treatment. (**D**) 6-d-old Col-0 and *jar1-1* seedlings grown vertically on MS medium containing nalacin, IAA, or DMSO as the mock control. Scale bar = 10 mm. (**E-F**) Quantification of the primary root length and hypocotyl length of seedlings shown in (**D**). Values represent means and ±SD (n ≥ 15). Statistical significances were analyzed by two-way ANOVA along with Tukey’s comparison test (\*\**P* < 0.01). Each data was derived from experiments that were performed three times with similar results, and representative data from one repetition were shown.

**Figure S6.**
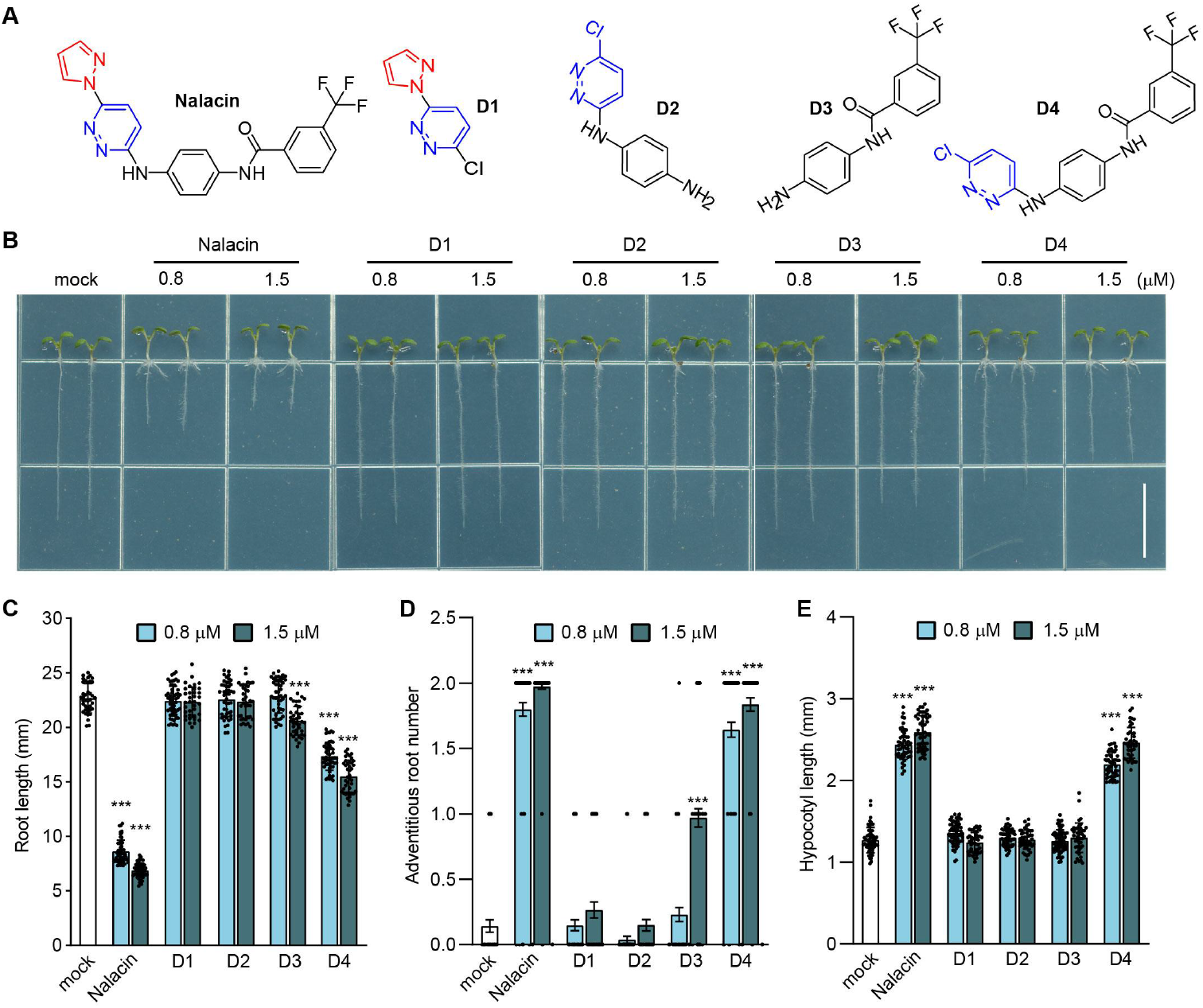
Structure-activity relationship analysis of nalacin. (**A**) Structure of nalacin and its derivatives D1, D2, D3, and D4. (**B**) Representative Col-0 seedlings vertically grown on MS medium containing chemicals for 6 days. Scale bar = 10 mm. (**C-E**) Quantification of the primary root length, adventitious root number and hypocotyl length of the seedlings treated by chemicals or DMSO as the mock control. Statistical significance was analyzed by two-way ANOVA along with Tukey’s comparison test (\*\*\**P* < 0.001). Data in (**B-E**) were derived from experiments that were performed three times with similar results, and representative data from one replicate were shown.

**Figure S7.**
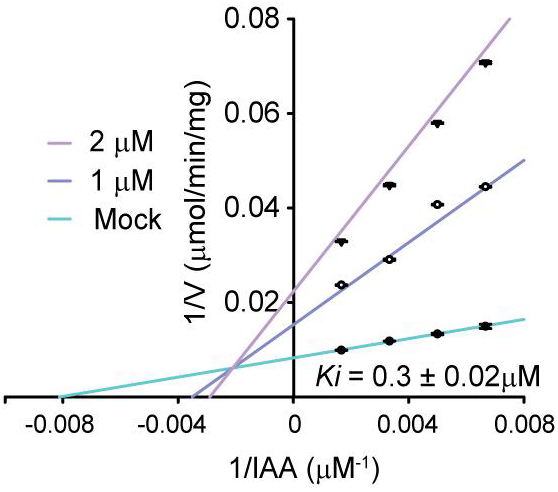
D4 inhibits the enzymatic activity of GH3.6. Kinetic analysis of the inhibition of D4 on GH3.6. Activity assays were performed with a fixed concertation of aspartate (10 mM) and gradient concentrations of IAA (0.15 mM −1 mM) in the absence or presence of D4. The production of IAA-Asp was detected by UPLC-MS.

**Figure S8.**
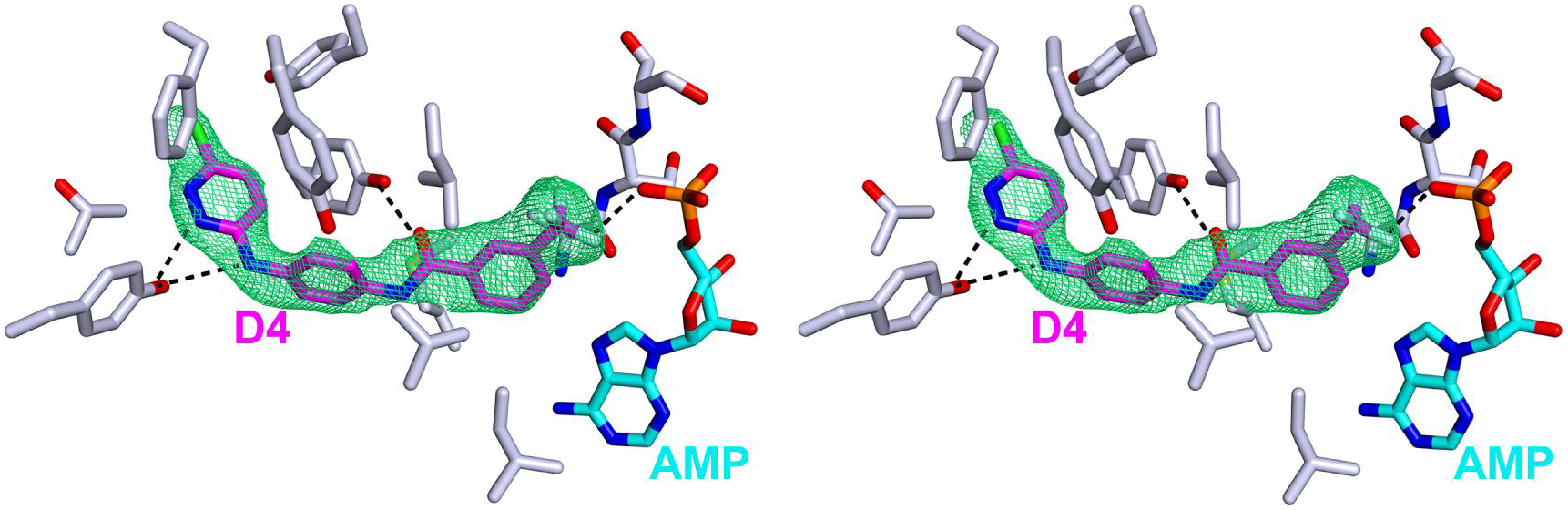
Stereo view of the density map of D4 in the GH3.6-AMP-D4 complex. Stereo view highlighting the density map of D4 in the GH3.6-AMP-D4 complex structure. The 2|Fo|–|Fc| s-weighted map is contoured at 1.0 σ.

**Figure S9.**
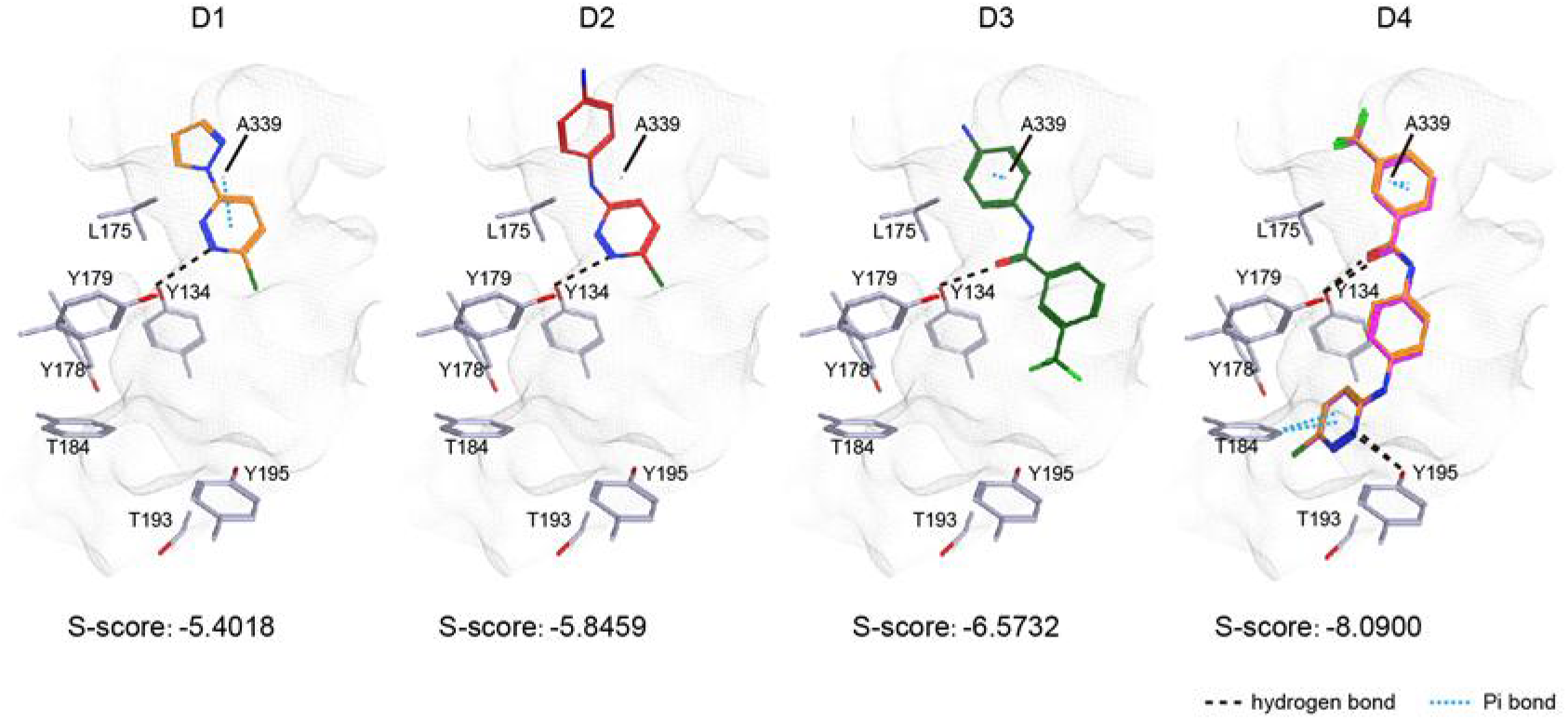
Site-views of GH3.6 in complex with molecular docked chemicals. D1, D2, D3, and D4 were docked to GH3.6 according to the procedure described in methods, and achieved s scores, the lower the score the stronger the binding affinity of the compound to the target protein. D1, D2, and D3 display different binding mode in the pocket of GH3.6. Molecular docked D4 (orange color) has almost the same binding pose with D4 from crystal structure (purple color).

**Figure S10.**
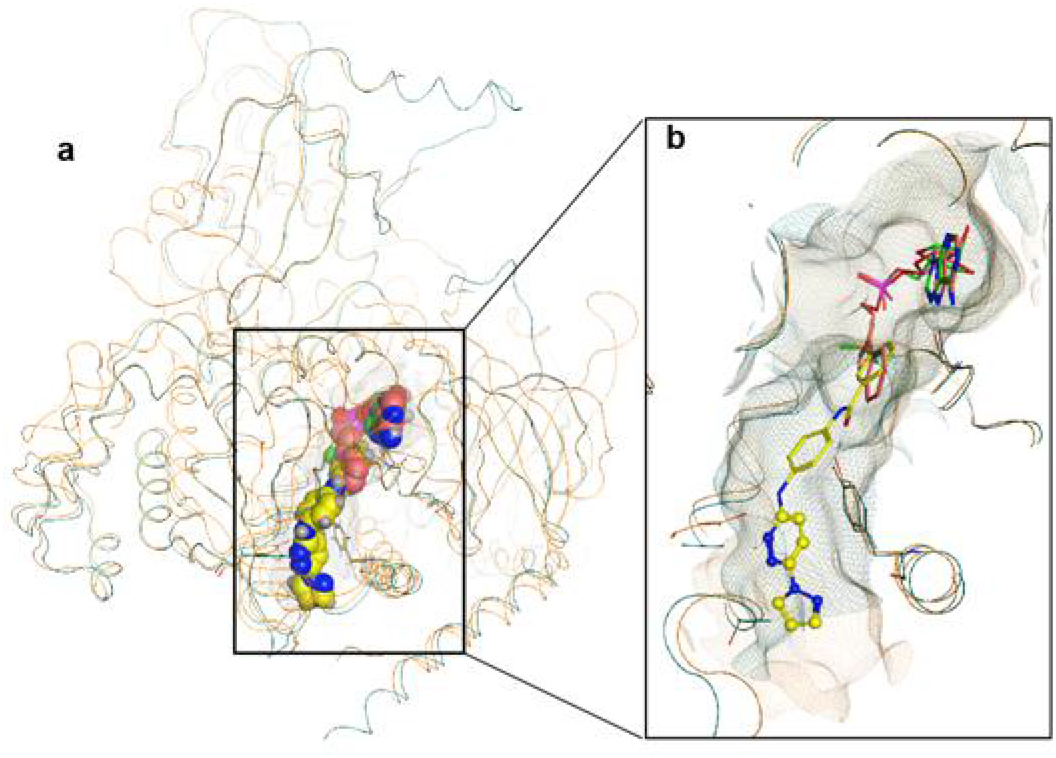
A pocket analysis of GH3s in complex with inhibitors. (**A**) The structure of GH3.6 (ribbon in green color) in complex with nalacin (spheres in yellow color) and AMP (spheres in green color) was superposed with the structure of VvGH3.1 (ribbon in orange color) in complex with AIEP (spheres in pink color) (PDB: 4B2G), a reaction intermediate mimic of IAA-AMP. (**B**) A site-view of GH3 pockets (GH3.6 in green mesh line; VvGH3.1 in orange mesh lines). The pyrazol-1-yl-pyrizadine moiety of nalacin (in ball-stick model) is in a deeper hydrophobic pocket of GH3.6, which could not directly participate the amino acid conjugation reaction.

**Figure S11.**
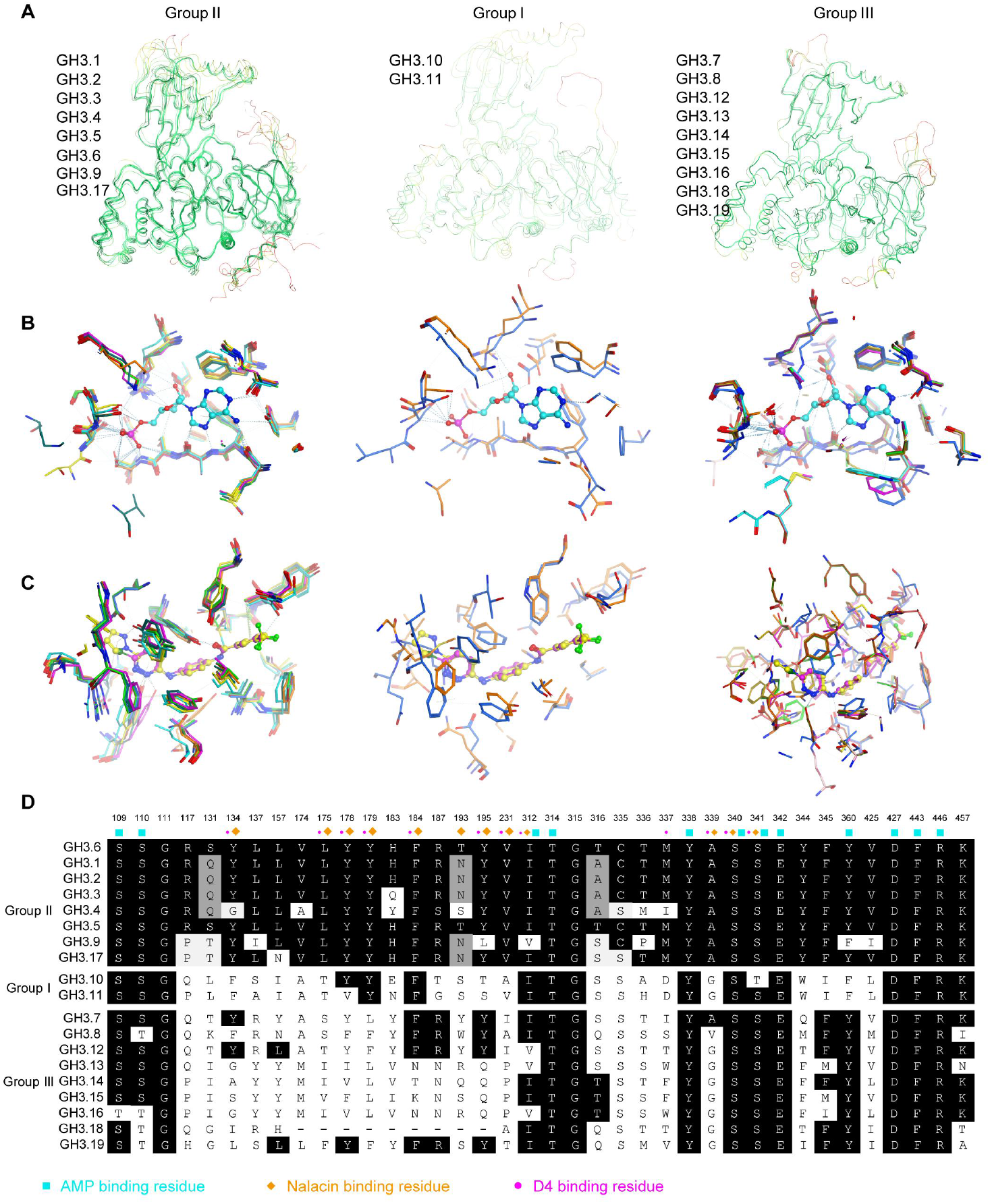
Structural analysis and comparison of distinctive groups of GH3s. The 3D structures of GH3s were downloaded from Protein Data Bank (5KOD for GH3.5, 4EPL for GH3.11, 4EPM for GH3.12) and AlphaFold Protein Structure Database (GH3.1, GH3.2, GH3.3, GH3.4, GH3.9, GH3.10, GH3.15, and GH3.17), or established by homology modelling with MOE (GH3.7, GH3.8, GH3.13, GH3.14, GH3.16, GH3.18, GH3.19), or originally resolved in this study (GH3.6 in complex with AMP and D4). (**A**) The superpose of overall fold of GH3s in group I, II, and III respectively. The ribbons are displayed in Root Mean Square Distance (RMSD) coloring, with low RMSD colored in green and increasing RMSD colored in yellow to red. (**B**) A site-view of the AMP binding sites. AMP is displayed in ball-stick model with cyan color. (**C**) A site-view of the binding sites for nalacin and D4. nalacin and D4 are displayed in ball-stick with yellow and purple colors, respectively. (**d**) Sequence alignment of residues that form the pocket of GH3s. Residues that interact with AMP, nalacin, and D4 are indicated with different symbols.

**Figure S12.**
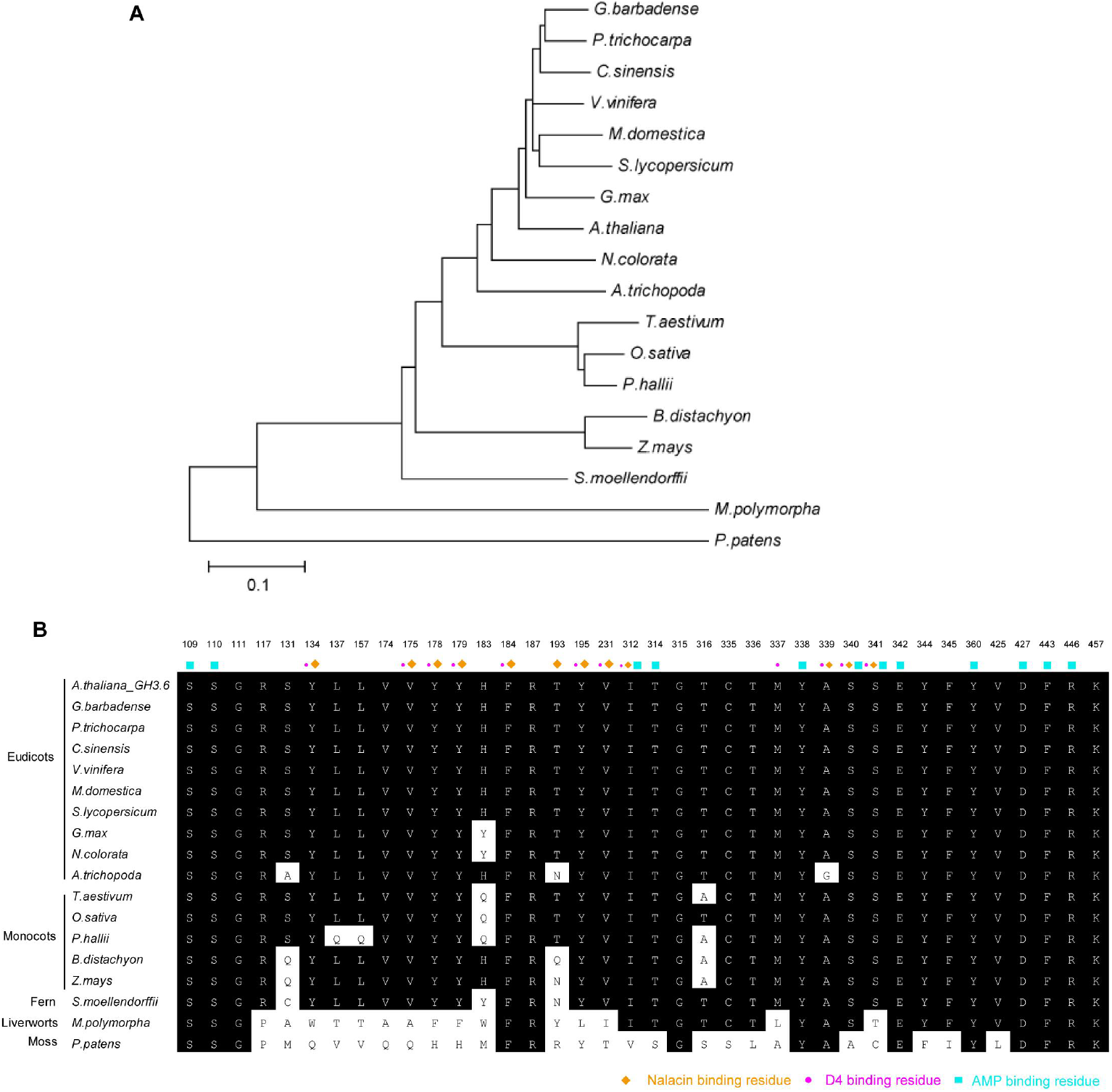
Phylogenetic analysis of GH3.6 proteins in various plant species. (**A**) The phylogeny tree of GH3.6 proteins in various species. (**B**) Sequence alignment of residues that form the pocket of group II GH3s in various species. Residues that interact with nalacin, D4, and AMP are indicated with different symbols.

**Figure S13.**
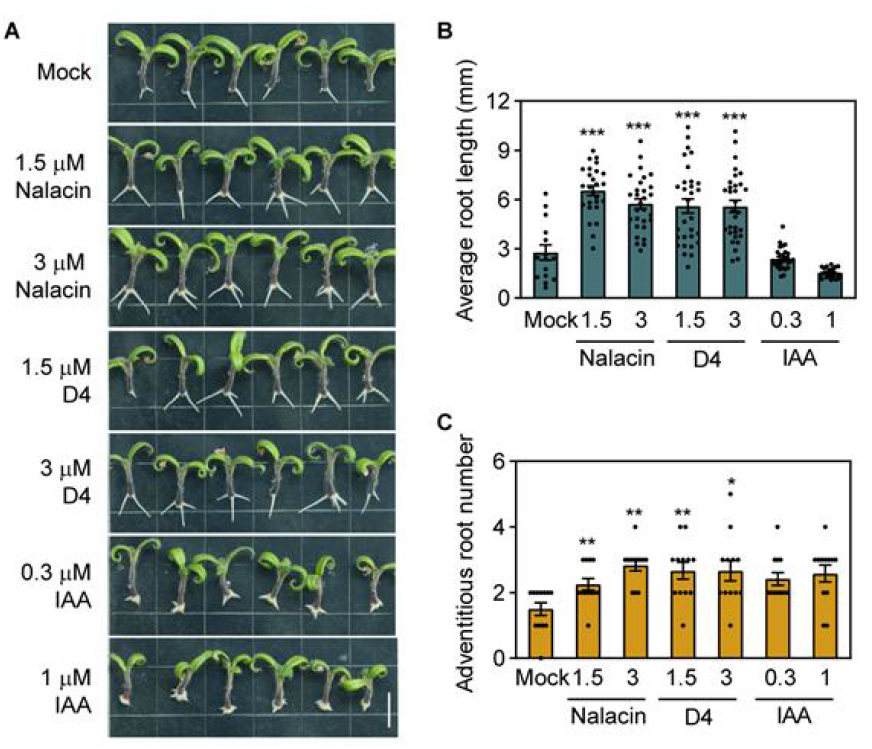
Nalacin and D4 promote adventitious root formation in tomato. **(A)** The roots of 7-day-old tomato seedlings were removed and transferred to MS medium containing nalacin, D4, IAA, or DMSO followed by additional cultivation for 6 days. Scale bar = 10 mm. **(B-C)** Quantification of the root length and adventitious root number shown in (**A**).

**Figure S14.**
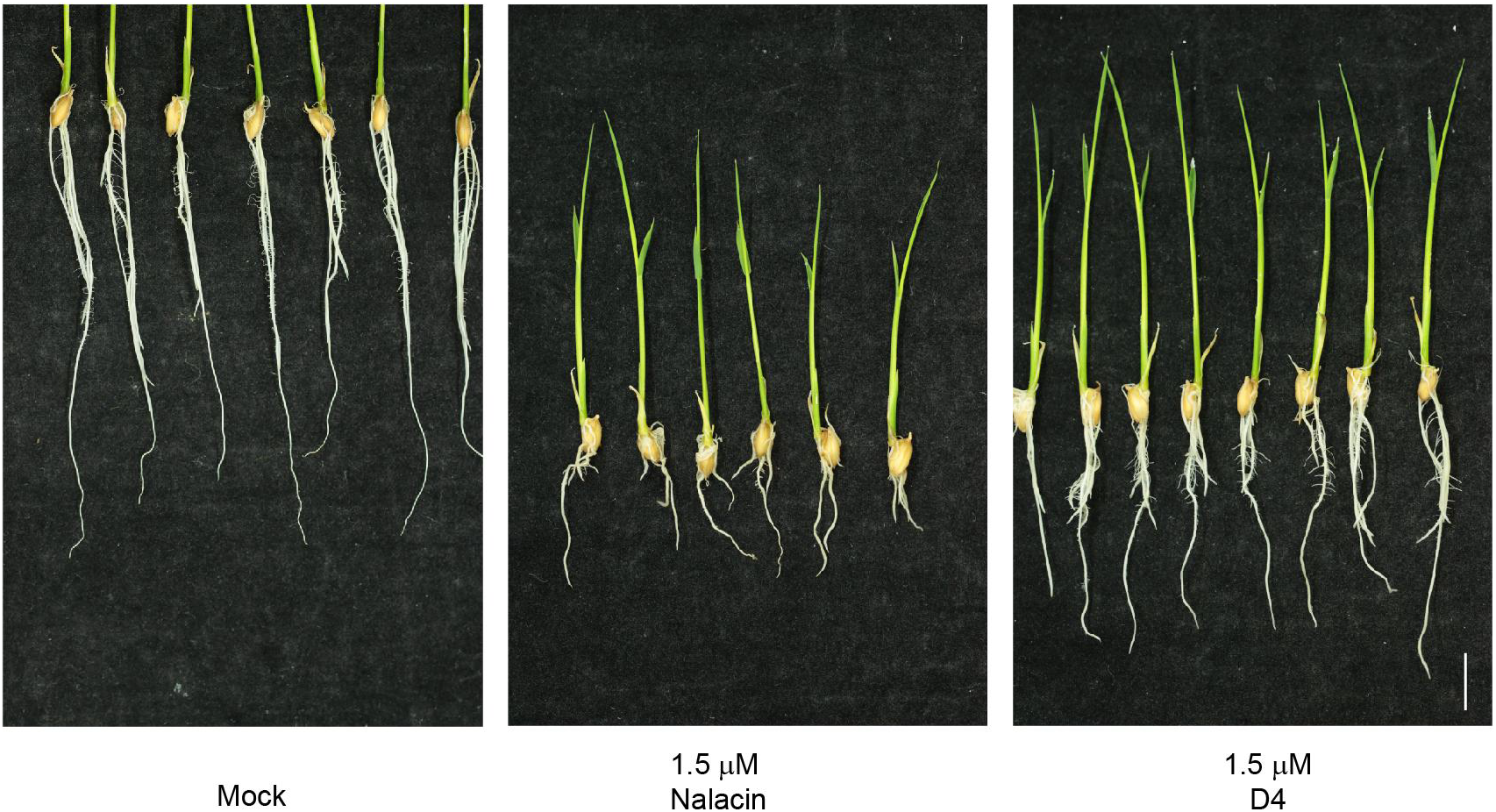
Nalacin and D4 regulate the root development of rice. The germinated seeds of rice (Nipponbare) were treated with nalacin, D4, or DMSO for 10 days in hydroponic solution. Scale bar = 1 cm.

**Supplementary Table 1.** Data collection and refinement statistics

| AtGH3.6-AMP-D4<br>PDB ID: 7VKA |  |
| --- | --- |
| **Data collection** |  |
| Space group | *P*6<sub>4</sub> |
| Cell dimensions |  |
| *a*, *b*, *c* (Å) | 205.9, 205.9, 65.7 |
| α, β, γ (°) | 90, 90, 120 |
| Resolution (Å) | 40-2.40 (2.49-2.40)<sup>a</sup> |
| *R*<sub>pim</sub> (%) | 5.2 (23.6) |
| *I* / *σ* | 6.3 (1.3) |
| Completeness (%) | 99.4 (96.4) |
| Redundancy | 11.0 (4.0) |
| No. of reflections<br>(total/unique) | 682,118/62,068 |
| **Refinement** |  |
| Resolution (Å) | 35.66-2.40 |
| Completeness (%) | 99.1 |
| No. of reflections | 61,910 |
| *R*<sub>work</sub> / *R*<sub>free</sub> (%) | 17.2/24.4 |
| No. atoms |  |
| Protein | 9,387 |
| AMP | 46 |
| GOL | 6 |
| D4 | 54 |
| Water | 437 |
| B-factors |  |
| Protein | 38.5 |
| AMP | 29.4 |
| GOL | 37.0 |
| D4 | 66.1 |
| Water | 35.5 |
| R.m.s deviations |  |
| Bond lengths | 0.008 |
| Bond angles (°) | 0.977 |

<sup>a</sup>Values in parentheses are for highest-resolution shell. One crystal was used for each data set.

