## Supplemental materials and methods for "Chemical genetic screening identifies nalacin as an inhibitor of GH3 amido synthetase for auxin conjugation"

### *SI Materials and Methods*

#### Plant materials and growth conditions

All Arabidopsis mutants and transgenic lines employed in this study are in the Col-0 background unless stated otherwise. *DR5::GUS* (1), *DR5::GFP* (2), *DII-VENUS* (3), △*8gh3* (4) and *axr2-1* (5) lines were described previously. Seeds were surface-sterilized with 75% ethanol and 0.1% Triton X-100 for 15 min and washed with 100% ethanol once, then cold-treated for 3 d at 4°C for imbibition in water, and germinated on half-strength Murashige and Skoog (MS) medium plates (2.22 g/L MS salts, 15 g/L sucrose, 0.5 g/L MES, and 5g/L gellan gum, pH 5.7 to 5.8) supplemented with the indicated concentrations of nalacin, IAA, Kyn, and PPBo. For light-grown seedlings, the plates were placed at 22°C under long-day photoperiods (16h: 8h, light: dark) for 6 days. In transient treatment assays, the 6-d-old seedlings were transferred into liquid MS medium supplemented with indicated compounds in 6-well plates and then incubated under light for various time intervals.

The tomato wild-type (*Solanum lycopersicum* cv. Micro-Tom) and *DR5::GUS* transgenic seeds (6) were sterilized with 75% ethanol followed by 4% (v/v) hypochlorite, then rinsed three times with sterile water. These seeds were germinated in water in 50 ml centrifuge tubes by shaking horizontally at room temperature. The germinated seeds were grown on half-strength MS medium (2.22 g/L MS salts, 15 g/L sucrose, 5 g/L gellan gum, pH 5.7 to 5.8) for 7 d, then seedlings with excised roots were transferred to MS media supplemented with indicated chemicals or DMSO as the mock control for 6 d. All the tomato plants were placed at 22°C with a 16 h-light/8 h-dark illumination cycle.

Rice (*Oryza sativa* cv. Nipponbare) seeds were soaked and incubated in water and 28°C for two days, then the germinated seedlings were transferred into Hoagland nutrient solution supplemented with indicated chemicals or DMSO as the mock control for 10 d. All seedlings were grown at 22°C with a 16 h-light/8 h-dark illumination cycle.

**Small molecule library and screen information**

A small chemical library of synthetic molecules of 11,800 structurally diverse chemicals was purchased from Life Chemical (https://lifechemicals.com/). The chemicals were dissolved by DMSO to produce 10 mM mother stock solutions. The chemicals were distributed by an automated robot (INTEGRA Assist Plus) into 24-well plates and mixed with half-strength MS medium (2.22 g/L MS salts, 15 g/L sucrose, 5 g/L gellan gum, pH 5.7 to 5.8), producing a solid screening medium with final concentration of 10 μM chemicals. Arabidopsis seeds (Col-0) were surface-sterilized followed by imbibition for 3 d at 4°C. Six to ten seeds were placed on screening medium and cultivated vertically for 5 days. The screen was performed by observing the changes in primary root lengths.

**Root hair length measurement and fluorescence observation**

The root hairs at maturation zones were photographed by a commercial seedling phenotyping platform (Dynaplant; http://www.yph-bio.com/DynaPlant.asp). For each plant, 10 individual roots were examined, and 80 root hairs were measured finally (n = 10 roots). The fluorescence in the primary root of *DR5::GFP* and *DII-VENUS* lines were observed by laser confocal microscopy (Zeiss LSM710). The excitation wavelengths were 488 nm and 514 nm along with 495-525 nm and 519-620 nm as the emission spectra for the fluorescence observation of GFP and YFP respectively.

**GUS staining**

Seedlings were grown on half-strength MS medium for 4 d, then transferred to the medium supplemented with indicated chemicals or DMSO as the mock control for additional treatment for 1 or 2 d. The seedlings were pre-washed with phosphate buffer saline and stained with GUS staining buffer (50 mM sodium phosphate buffer, pH 7.0, 10 mM $Na_2EDTA$, 0.5 mM $K_4[Fe(CN)_6]\cdot 3H_2O$,

0.5 mM $K_3[Fe(CN)_6]$, 0.1% Triton X-100, and 1 mg/mL X-Gluc) for optimal time at 37°C. 70% ethanol was used to terminate the reaction. The seedlings were mounted on slides in Hoyer's solution (chloral hydrate:water:glycerol; 8:3:1; w/v/v) and examined by microscopy.

**Measurement of IPA, IAA and IAA catabolites**

For quantification of Trp, IPA, IAA, oxIAA, IAA-Asp and IAA-Glu, 6-d-old seedlings of Col-0 were incubated in liquid MS medium supplemented with 3 μM nalacin or DMSO as the mock control for two hours. After treatment, the seedlings were washed with Milli-Q quickly and sampled in liquid nitrogen. The contents of these metabolites were determined by Wuhan Greenword Creation Technology Company (http://www.greenswordcreation.com/index.html) with UPLC-MS/MS (Thermo Scientific TSQ Quantiva-Stage Quandrupole Mass Spectrometer). In brief, about 50 mg of fresh leaves was ground in liquid nitrogen into a powder and then extracted with 1.0 mL 80% cold methanol (v/v) at 4°C for 12 h. After centrifugation at 4°C at 12,000 rpm for 10 min, the supernatant was collected and evaporated with mild nitrogen stream at 35°C followed by re-dissolving in 100 μL $H_2O$/ acetonitrile (90/10, v/v) for loading samples. Gradient elution was performed with solvent A (water with 0.1% formic acid) and solvent B (acetonitrile). The effluent was introduced into mass spectrometer with the optimal setting as follow: positive ionization mode; capillary voltage, 3000 V; sheath gas flow, 50 Arb; auxiliary gas flow, 12 Arb; sweep gas flow, 4 Arb; collision gas, 1.5 mTorr; ion transfer tube temperature, 350 °C; vaporizer temperature, 300 °C; spray voltage, 3000 V. Three independent replicates were used for each treatment. Data were analyzed and processed using Thermo Scientific Xcalibur 2.1 data system (Thermo Scientific, USA).

**GH3 enzymatic activity assay and kinetic analysis**

AtGH3.3, AtGH3.6, and AtGH3.17 were expressed in *E. coli* BL21 (Rosetta) using pDEST17 vector with an N-terminal His-tag. Cells were cultured at 37°C in LB medium to an $OD_{600}$ for 3 h, then these proteins were induced by 0.5 mM isopropyl-β-D-thiogalactopyranoside overling at 18°C. Cells were collected by centrifugation and resuspended in lysis buffer (50 mM Tris-HCl, pH 8.0, 500 mM NaCl, 20 mM imidazole, 1 mM DTT, 1 mM PMSF). Protein purification was performed using the ÄKTA pure system (GE Healthcare) according to the manufacturer's instructions. Purified GH3 proteins were concentrated and flash-frozen with liquid nitrogen, then stored at - 80°C. IAA-amido synthetase assay included 50 mM Tris-HCl (pH 8.0), 2 mM $MgCl_2$, 2 mM ATP, 1 mM DTT, 10 mM Asp/Glu, 0.15 mM ATP, concentration gradients IAA (200–1000 μM) and indicated nalacin or DMSO at 25°C for 30 min. The production of IAA-Asp or IAA-Glu were monitored by UPLC-MS. The enzyme kinetics and the inhibitory mode of nalacin were analyzed with Sigmaplot software (version 14).

**Biolayer interferometry analysis of binding activity of nalacin with GH3 proteins**

Biolayer interferometry assay were conducted as described previously (7) by Octet® R2 protein analysis system (Sartorius). Briefly, purified GH3 proteins were labelled with a 1:1 molar ratio of biotin to protein, followed by removal of free biotin with a gravitational desalination column (#G-MM-ITG, www.bomeida.con). Super streptavidin biosensors (SSA) with or without labelled GH3s proteins washed with PBS buffer. Dilution gradients of nalacin were generated by PBS (0.5% DMSO) buffer. BLI measuring protocol for each step as followed: baseline in PBS buffer for 60 s, association in indicated nalacin solution for 60 s, dissociation in PBS buffer for 60 s, and wash sensors in PBS buffer for 300s. When all the concentration of nalacin were detected, the global fitting method were applied to generate kinetic affinity constants of GH3s with nalacin, the K*d* values of GH3.6 with nalacin and AMP/ATP were analyzed by steady-state affinity. The data were analyzed by Fortebio data analysis software.

**Crystallization, X-ray data collection and structure determination**

The full-length Arabidopsis *GH3.6* (*GH3.6*) gene was isolated from an Arabidopsis cDNA library and cloned into the first multiple cloning site of a modified RSFDuet-1 vector (Novagen) with an N-terminal $His_6$-SUMO tag. For expression of GH3.6, home-made Rosetta (DE3) competent cells were transformed with the corresponding vector and then cultured using Luria-Bertani (LB) medium supplemented with proper antibiotics at 37 °C to an $OD_{600}$ of 1.0-1.2. Then, GH3.6 expression was induced with 0.2 mM isopropyl β-D-1-thiogalactopyranoside (IPTG), and cells were further incubated overnight at 18 °C. The expressed GH3.6 protein was first purified using Ni-sepharose 6 affinity beads (GE Healthcare). After removal of the N-terinmal $His_6$-SUMO tag with Ulp1 (SUMO protease; home-made), the resulting AtGH3.6 protein was further purified on a HiLoad 16/600 Superdex 200 column (GE Healthcare) equilibrated in a buffer of 20 mM Tris-HCl, pH 7.5, 100 mM NaCl. The pure fractions were pooled together and concentrated, then kept at -80 °C until further use.

For crystallization of the GH3.6-AMP-D4 complex, 468 μL of GH3.6 at a concentration of 7.5 mg/mL (diluted from higher concentration of stock solution kept at -80 °C) in a buffer of 20 mM Tris-HCl, pH 7.5, 100 mM NaCl; then 5 mM AMP (final concentration; 2.5 μL of 1 M stock solution), 5 mM TCEP (final concentration; 5 μL of 0.5 M stock solution) and 25 μL of D4 molecules (20 mM stock solution in DMSO) were added; and the resulting mixture was incubated at 20 °C for 2-3 hrs. Then, the protein sample was centrifuged for 10 mins (13,000 rpm at 6 °C), and the supernatant was taken for crystallization trials.

All the crystallization trials were done by using the sitting-drop vapor-diffusion method at 20 °C. Crystals of the GH3.6-AMP-D4 complex were grown in 0.2 M Magnesium chloride, 0.1 M Tris pH 8.5, 20% PEG 8000. Then, all crystals were stabilized in mother liquor containing 20% glycerol and flash frozen in liquid nitrogen.

All X-ray diffraction data were collected at a wavelength of 0.979 Å on beamline 19U1 of the Shanghai Synchrotron Radiation Facility (SSRF) (8). The diffraction data sets were processed with HKL3000 (9). The structure was solved by molecular replacement in PHASER (10) with the structure of GH3.5 (PDB: 5KOD) (11) as a search model. Then the model was manually built and refined using Coot (12) and refined with PHENIX (13). The final model of the GH3.6-AMP-D4 complex was refined to 2.40 Å. Supplementary table 1 summarizes all the statistics for data collection and structural refinement. Structural figures were prepared using PyMOL (The PyMOL Molecular Graphics System, Schrödinger).

**Molecular docking simulation**

Docking simulation was performed using the molecular modelling program Molecular Operating Environment (MOE) (Chemical Computing Group, 2020.09). The chain B of GH3.6-AMP-D4 (originally dissolved in this study with resolution: 2.40 Å) was selected for docking and prepared using the QuickPrep function module for fixation and protonation. The docking site of was defined as D4 binding site. The candidate molecules including nalacin, D1, D2, D3, and D4 were prepared by Wash module and Conformational Search before submitting to the docking simulation. For the docking process, 100 output poses per molecule were generated and and docked to the binding site with Induced Fit refinement. The final interaction energy between GH3.6 and candidate molecules were evaluated by Generalized Born solvation model (GBVI), a forcefield-based scoring function that estimates the binding free energy of binding of the ligand from a given pose. One pose per molecule with best S score was output and displayed graphically.

**Homology modelling of group III GH3s**

Homology modelling was performed using the molecular modelling program MOE. The 3D structure file of GH3.12 (PDB: 4EPM, Resolution: 2.10 Å), as a member of group III GH3, was downloaded from the Protein Data Bank (https://www.rcsb.org/), and the chain A of 4EPM was selected as template for homology modelling. We aligned the sequences of GH3.7, GH3.8, GH3.13, GH3.14, GH3.16, GH3.18, and GH3.19 using the Alignment module, and built three-dimensional structures using Homology model module. The final output model is the one with the best-scoring of the electrostatic solvation energy, which is calculated using a Generalized Born/Volume Integral (GBVI) methodology (14).

**Chemical preparation**

Nalalcin and D1-4 were synthesized by The Chemical Biology Laboratory at The University of Tokyo according to the procedures shown in below. The chemicals were confirmed by $^{1}$H NMR and $^{13}$C NMR. Electrospray ionization high resolution mass spectrometry (ESI-HRMS): *m/z* calculated for $C_{21}H_{16}F_3N_6O$ (nalacin) $[M+H]^+$: 425.1338, found 425.1324; *m/z* calculated for $C_7H_6ClN_4$ (D1) $[M+H]^+$: 181.0281, found 181.0274; *m/z* calculated for $C_{10}H_9ClN_4$ (D2) $[M+H]^+$: 221.0594, found 221.0582; *m/z* calculated for $C_{14}H_{12}F_3N_2O$ (D3) $[M+H]^+$: 281.0902, found 281.0891; *m/z* calculated for $C_{18}H_{13}ClF_3N_4O$ (D4) $[M+H]^+$: 393.0730, found 393.0719.

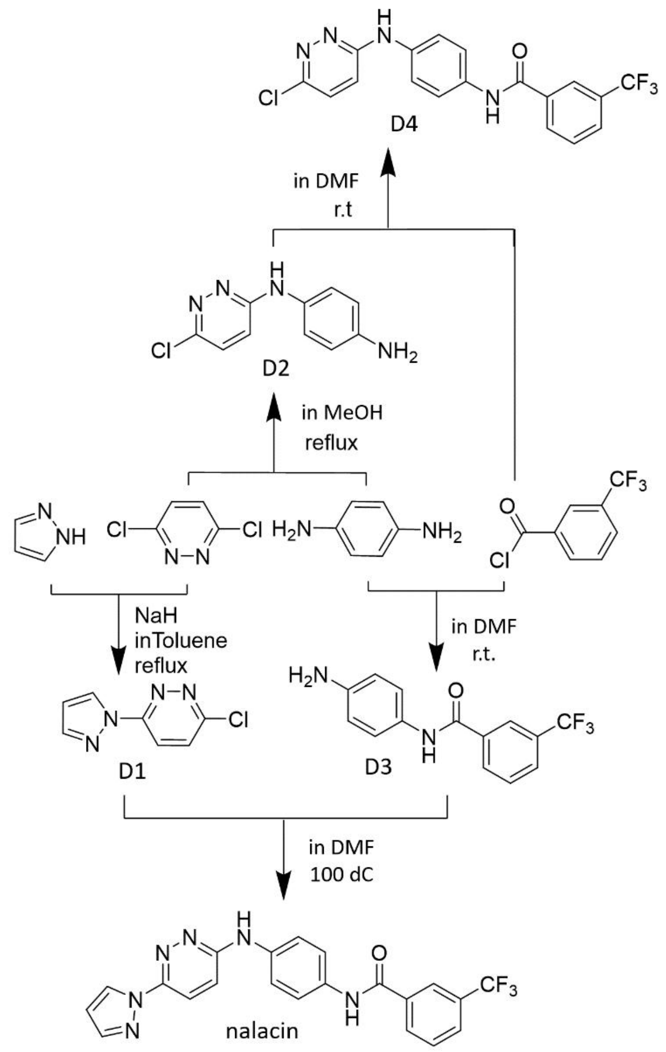


A schematic graph of chemical synthesis pathways for nalacin and D1-4

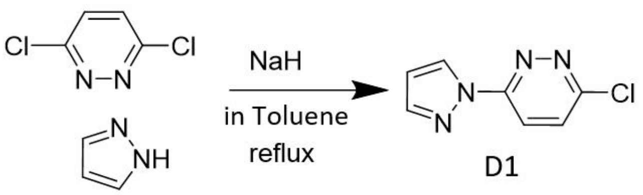


A schematic graph of chemical synthesis pathway for D1

To a solution of 3.6-dichloropyridazine (2.4 g, 16.2 mmol) and pyrazole(400 mg, 5.9 mmol) in toluene (20 mL), NaH (60％,269.3 mg, 6.7 mmol) was added slowly. Then the reaction mixture was refluxed for 1 h. Ethyl acetate (200 mL) and a small portion of MeOH were added to the reaction solution, which was then washed with water and brine. The organic layer was dried over $Na_2SO_4$ and the solvent was evaporated. The residue was recrystallized (hexane/ethyl acetate) to give 637.1 g of white solid 60%) identified as D1.

$^1$H-NMR (500 MHz, $CDCl_3$) δ: 8.72 (d, J = 3.0 Hz, 1H), 8.21 (d, J = 9.5 Hz, 2H), 7.80 (s, 1H), 7.63 (d, J = 9.5 Hz, 2H), 6.55 (d, J = 3.0 Hz, 1H). $^{13}$C-NMR (126 MHz, $CDCl_3$) δ: 154.35, 153.74, 143.41, 130.51, 127.54, 120.02, 109.30. ESI-HRMS: *m/z* calculated for $C_7H_6ClN_4$ $[M+H]^+$: 181.0281, found 181.0274.

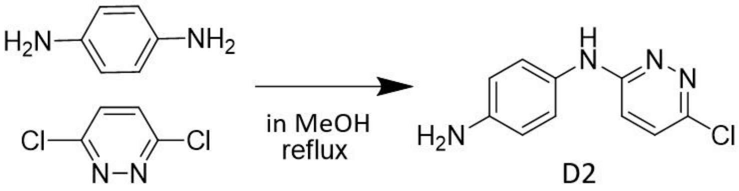


A schematic graph of chemical synthesis pathway for D2

A solution of 1.4-phenylenediamine (432 mg, 4.0 mmol) and 3.6-dichloropyridazine (595 mg, 4.0 mmol) in 20 mL of MeOH was refluxed overnight. Subsequently, the solution was cooled to room temperature and the solvent was evaporated under reduced pressure. The residue was purified by column chromatography (silica gel, hexane/ethyl acetate) to give 0.4 g of white solid (45%) identified as D2.

$^1$H-NMR (500 MHz, $CDCl_3$) δ: 7.16 (d, J = 9.5 Hz, 1H), 7.06 (d, J = 8.5 Hz, 1H), 6.97 (br, 1H), 6.84 (d, J = 9.5 Hz, 1H), 6.70 (d, J = 8.5 Hz, 2H), 3.72 (br, 2H). $^{13}$C-NMR (126 MHz, $CDCl_3$) δ: 158.93, 147.23, 144.63, 129.20, 129.03, 125.83, 115.97, 115.17. ESI-HRMS: *m/z* calculated for $C_{10}H_9ClN_4$ $[M+H]^+$: 221.0594, found 221.0582.

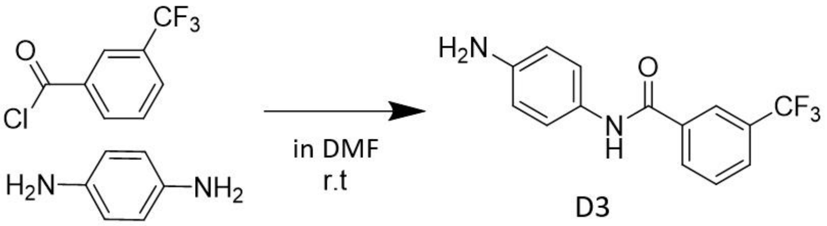


A schematic graph of chemical synthesis pathway for D3

To an ice-cooled solution of 1.4-phenylenediamine (3.29 g, 30.43 mmol) in DMF (100 mL), 3-(trifluoromethyl)benzoyl chloride (2.12 g, 10.14 mmol) was added dropwise. Then reaction mixture was stirred for 3 h at room temperature. Ethyl acetate (200 mL) was added to the reaction solution, which was then washed with water and brine. The organic layer was dried over $Na_2SO_4$ and the solvent was evaporated. The residue was purified by column chromatography (silica gel, hexane/ethyl acetate) to give 2.21 g of yellow oil (78%) identified as D3.

$^1$H-NMR (500 MHz, $CDCl_3$) δ: 8.34 (br, 1H), 8.08 (s, 1H), 7.99 (d, J = 7.5 Hz, 1H), 7.72 (d, J = 7.5 Hz, 1H), 7.51 (t, J = 7.5 Hz, 1H), 7.34 (d, J = 8.0 Hz, 2H), 6.61 (d, J = 8.0 Hz, 2H), 3.66 (br, 2H). $^{13}$C-NMR (126 MHz, $CDCl_3$) δ:164.39, 143.83, 135.90, 130.89 (q, J = 33.0 Hz), 130.31, 129.09, 128.69, 127.89 (q, J = 3.7 Hz), 124.02 (q, J = 3.7 Hz), 123.65 (q, J = 272.4 Hz), 122.67, 115.25. ESI-HRMS: *m/z* calculated for $C_{14}H_{12}F_3N_2O$ $[M+H]^+$: 281.0902, found 281.0891.

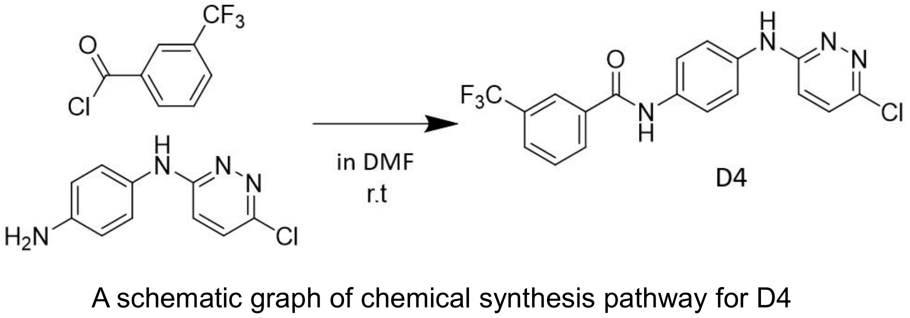


A schematic graph of chemical synthesis pathway for D4

To an ice-cooled solution of D2 (216 mg, 0.77 mmol) in DMF (7.7 mL), 3-(Trifluoromethyl)benzoyl Chloride (241 mg, 1.16 mmol) was added dropwise. Then reaction mixture was stirred for 3 h at room temperature. Ethyl acetate (200 mL) was added to the reaction solution, which was then washed with water and brine. The organic layer was dried over $Na_2SO_4$ and the solvent was evaporated. The residue was purified by column chromatography (silica gel, hexane/ethyl acetate) to give 0.195 g of yellow solid (64%) identified as D4.

$^1$H-NMR (500 MHz, DMSO) δ: 10.44 (s, 1H), 9.49 (s, 1H), 8.31 (s, 1H), 8.28 (d, J = 8.0 Hz, 1H), 7.97 (d, J = 8.0 Hz, 1H), 7.79 (d, J = 8.0 Hz, 1H), 7.75 (d, J = 9.0 Hz, 2H), 7.71 (d, J = 9.0 Hz, 2H), 7.56 (d, J = 9.0 Hz, 1H), 7.20 (d, J = 9.0 Hz, 1H). $^{13}$C-NMR (126 MHz, DMSO) δ: 163.60, 156.74, 147.11, 1236.37, 135.84, 133.04, 131.73, 129.67, 129.14 ( q, J = 31.9 Hz), 127.98 ( q, J = 3.3 Hz), 124.15 (q, J = 3.9 Hz,),124.00 (q, J=354.02). ESI-HRMS: *m/z* calculated for $C_{18}H_{13}ClF_3N_4O$ $[M+H]^+$: 393.0730, found 393.0719.

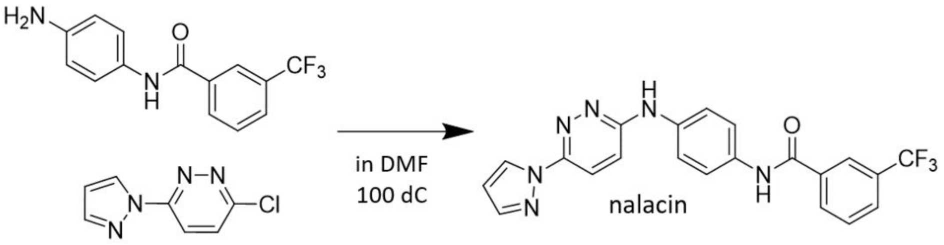


A schematic graph of chemical synthesis pathway for nalacin

The mixture of D3 (1.5 g, 5.5 mmol) and D1 (0.995 g, 5.5 mmol) in MeOH (5.5 mL) was refluxed for a week at 100 °C. Dichloromethane (50 mL) was added to the reaction mixture, which was then washed with water and brine. The organic layer was dried over $Na_2SO_4$ and the solvent was evaporated. The residue was purified by column chromatography (silica gel, hexane/ethyl acetate) to give solid, which was then recrystallized to give 1.1 g of pale-yellow solid (47%) finally identified as nalacin.

$^1$H-NMR (500 MHz, $CDCl_3$) δ: 10.45 (s, 1H), 9.51 (s, 1H), 8.69 (d, J = 3.0 Hz, 1H), 8.33 (s, 1H), 8.30 (d, J = 8.0 Hz, 1H), 8.05 (d, J = 9.5 Hz, 1H), 7.97 (d, J = 8.0 Hz, 1H), 7.86 (s, 1H), 7.77-7.82 (m, 5H), 7.38 (d, J = 9.5 Hz, 1H), 6.62 (d, J = 3.0 Hz, 1H). $^{13}$C-NMR (126 MHz, DMSO) δ: 163.56, 156.56, 148.80, 141.81, 1236.89, 135.87, 132.67, 131.73, 129.67, 129.15 (q, J = 32.2 Hz), 127.96 (q, J = 3.6 Hz), 124.15 (q, J = 3.8 Hz), 124.00 (q, J = 272.5 Hz), 121.30, 120.10, 119.72, 118.77, 118.50, 108.32. ESI-HRMS: *m/z* calculated for $C_{21}H_{16}F_3N_6O$ $[M+H]^+$: 425.1338, found 425.1324.
